# Relational Structure Constrains Individual Value Estimates in Visual Working Memory

**DOI:** 10.64898/2026.07.27.740746

**Authors:** Jaeseob Lim, Sang-Hun Lee

## Abstract

Reflecting how we organize visual experience in everyday memory, visual working memory is increasingly understood as a system in which individual item representations are organized within structures rather than maintained in isolation. Among these, relational structure may be especially consequential because, by specifying how one item value lies relative to another within a feature space, it could allow information about one remembered value to constrain which values are plausible for the other. Yet demonstrating such constraint is challenging because item-specific mnemonic evidence and relational evidence ordinarily support essentially the same estimate. We broke this equivalence with biased post-encoding feedback for one item, making item-specific and relation-based predictions for the other diverge. Across three experiments, participants remembered two sequentially presented orientations, with feedback for one shifted slightly clockwise or counterclockwise from its actual value. Participants incorporated this bias into memory for the feedback-provided orientation; critically, it also appeared in reports of the other orientation, which received no feedback, in the direction predicted by the signed angular offset linking the two remembered values. This feedback transfer weakened with increasing angular separation but occurred in both directions between the first and second orientations. These findings show that relational structure directly constrains individual value estimates in visual working memory, even for items encountered separately. By dissociating normally coincident item-specific and relation-based predictions, our approach reveals an otherwise hidden relational contribution. A probabilistic account explains these findings through joint inference from uncertain item-specific and relational evidence, with their relative uncertainties governing transfer strength.

## Introduction

Working memory (WM) is often studied as a limited-capacity system for maintaining information no longer available to perception (Bays & Husain, 2008; Luck & Vogel, 1997). However, its role likely extends beyond maintaining item information in isolation. In everyday cognition, sensory inputs are often fragmented, fleeting, or incomplete, creating a need for WM to organize them into structured representations that support flexible inference. This broader view aligns with longstanding accounts in terms of schemas, chunks, frames, and relational structures, as well as with more recent views of WM as a compositional workspace in which information is selected, bound, and transformed to meet current goals (Bartlett, 1995; D’Esposito & Postle, 2015; Gentner, 1983; Halford et al., 2010; Oberauer, 2009; Wyble et al., 2025).

One well-studied form of such structure in visual WM is the use of summary representations of multiple items. Observers efficiently extract ensemble statistics—such as mean size, orientation, location, or expression—from visual displays (Alvarez & Oliva, 2009; Ariely, 2001; Chong & Treisman, 2003; Haberman & Whitney, 2007; Parkes et al., 2001). These summaries can shape visual WM: memory for individual items is biased toward group means or global display averages, suggesting that item-level information is integrated with higher-level summaries (Brady & Alvarez, 2011, 2015). Related models and empirical findings further show that grouping, chunking, and similarity-based clustering can organize multiple items into higher-order units that shape recall precision and bias (Brady & Tenenbaum, 2013; Corbett, 2017; Orhan & Jacobs, 2013; Son et al., 2020).

However, ensemble statistics provide only compressed information about structure—what is typical of a set—without specifying how individual elements relate to one another. Relations provide a different form of structure. By specifying how elements are linked within a scene, environment, or feature space, relational structure can be abstracted from the particular sensory contents that instantiate it, allowing the same relational form to organize new experiences and support flexible generalization (Kemp & Tenenbaum, 2008; Summerfield et al., 2020; Tenenbaum et al., 2011; Tervo et al., 2016). Everyday navigation highlights why relational structure is useful (Behrens et al., 2018). As we hike along winding trails, we encounter successive path segments, bends, and intersections rather than perceiving the entire route at once. Representing the relative turns linking these local encounters allows them to be integrated into a coherent route despite changes in viewpoint or body orientation. More generally, relational information and structure can organize separately encountered elements into a representation of how they fit together.

This raises a more specific question for visual WM: when the feature values of multiple items are encoded and maintained, does the relation between them merely describe how those values are organized, or does it become part of what constrains each remembered value? One possibility is that separate item representations are organized into an integrated relational structure in which individual item values are linked by relational information. In such a structure, an item’s feature value may be inferred primarily from its own item-specific mnemonic evidence while also being constrained by its relation to another item. Building on prior evidence for structured representations in visual WM (Brady & Alvarez, 2011; Orhan & Jacobs, 2013), this proposal focuses on the relational component of such structure (Bae & Luck, 2017). For example, remembering two orientations (first 30°, second 50°) may involve representing one item as approximately 30° and the other as approximately 50°, while also representing their signed relation: that the second item is approximately 20° counter-clockwise from the first. Critically, this relation is not external to the remembered values. It provides relational evidence about those values because it specifies how one value is positioned relative to another within the same feature dimension. In this sense, the relation is not merely a cue for otherwise separate item-value representations; it is part of what constrains the remembered values themselves.

Prior work has shown that relational or configurational information can shape visual WM performance, including cases in which observers are sensitive to whether relations among items are preserved or violated, or in which the spatial configuration of surrounding items affects memory for a cued target (Bateman et al., 2018; Dent, 2009; Jiang et al., 2000). These findings demonstrate that relations can influence memory judgments about individual items. However, because the relational information in these tasks often serves as a context for retrieval, comparison, or change detection, they do not establish whether a relation defined over item values directly constrains inference about an individual item’s feature value.

The key prediction follows from a contrast with influential item-based accounts of visual WM, which generally explain memory in terms of item-specific storage or evidence without explicitly positing relations that couple different item values (Bays & Husain, 2008; Luck & Vogel, 1997; Wilken & Ma, 2004; W. Zhang & Luck, 2008). Under such accounts, the remembered value of each orientation should depend primarily on its own item-specific evidence. If the two orientations are linked by a signed angular relation, however, one item’s value can also constrain inference about the other. For example, a memory representation of the second item may be constrained both by its own item-specific mnemonic evidence, favoring approximately 50°, and by a relation-based prediction derived from its signed angular relation to the first item, such as being approximately 20° counter-clockwise from it (**Fig. 1A**). The critical question is whether this relational evidence actually contributes to inference about the second item’s value.

**Figure 1.**
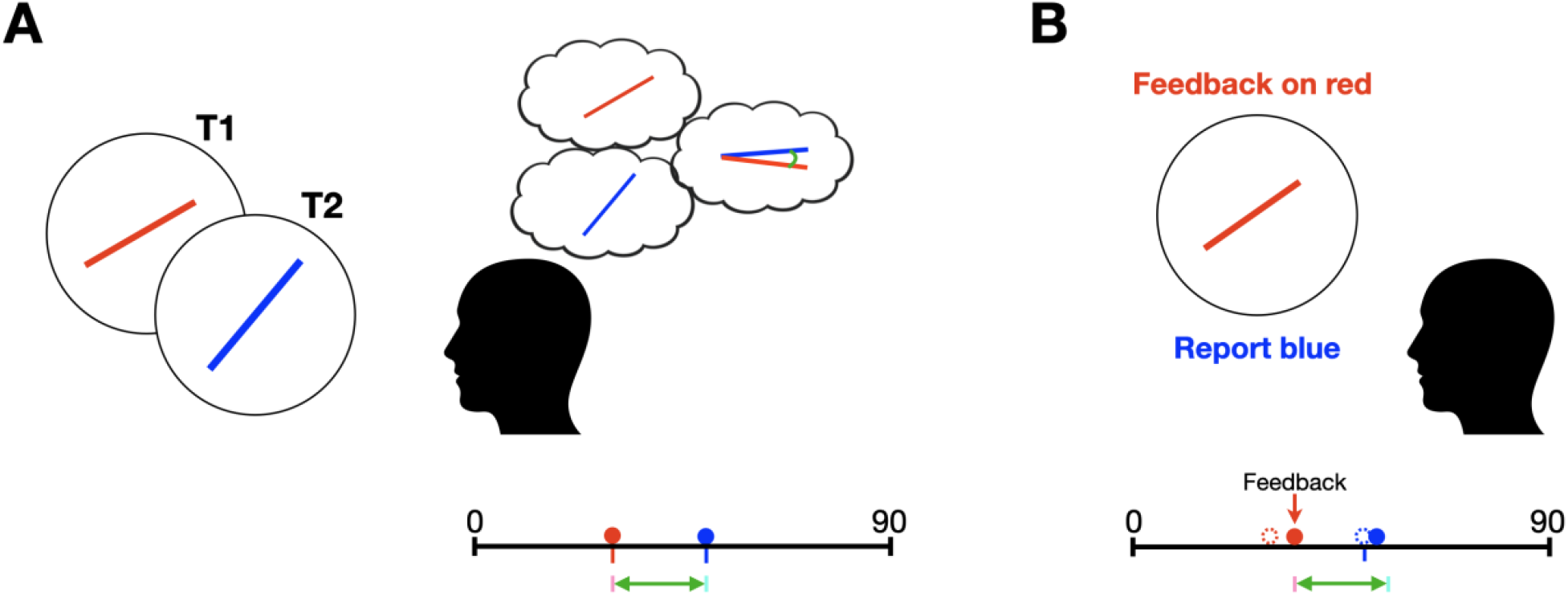
Relation-constrained item representations and selective feedback perturbation in visual WM. **(A)** A red orientation (T1) and a blue orientation (T2) are encoded sequentially and maintained in visual working memory. The separate red and blue thought bubbles depict item-specific mnemonic representations, whereas the joint thought bubble and green arrows depict the signed angular relation linking the two orientations. The black horizontal line represents orientation feature space; red and blue circles indicate memory estimates for each item, red and blue ticks indicate item-specific evidence, and cyan and pink ticks indicate relation-based predictions for the blue and red orientations, respectively. In an independent-item account, each memory estimate is determined by its own item-specific evidence without constraint from the other item. In a relation-constrained account, each estimate is additionally constrained by a relation-based prediction derived from the other orientation through their remembered signed relation (green arrow). **(B)** Feedback specifies a value for the red orientation that differs from its pre-feedback estimate. Dotted circles indicate the pre-feedback estimates, and solid circles indicate the feedback-specified value of the red orientation and the resulting estimate of the blue orientation. If the two orientations are represented independently, the estimate of the blue orientation should remain unchanged. If they are linked by their signed angular relation, however, the estimate of the blue orientation should shift in a relation-consistent direction, even though feedback was not provided for the blue orientation.

Under ordinary conditions, however, item-specific and relational contributions are difficult to disentangle because they typically support nearly the same value. Suppose mnemonic evidence for item A favors 30°, mnemonic evidence for item B favors 50°, and their remembered signed relation is +20°. The relation-based prediction for B—A+20°—is then also 50°, coinciding with the value supported by B’s own mnemonic evidence. A report near 50° therefore cannot reveal whether B was inferred solely from its item-specific evidence or was additionally constrained by its relation to A.

Selective post-encoding feedback breaks this equivalence by separating the values supported by the two sources. If biased feedback specifies a value of 35° for item A, the remembered relation now predicts 55° for item B—A+20°—whereas B’s own mnemonic evidence still favors approximately 50°(**Fig. 1B**). An independent-item account therefore predicts little systematic change in B’s report, whereas a relation-constrained account predicts a shift toward 55°. Because both sources are uncertain (Bays et al., 2024; Bays & Husain, 2008; Van den Berg et al., 2012; Wilken & Ma, 2004), the reported value should reflect a reliability-weighted compromise between item-specific and relational evidence, consistent with reconstructive accounts in which uncertain item-specific evidence is combined with other relevant information to determine remembered values (Brady & Alvarez, 2011; Hemmer & Steyvers, 2009; Huttenlocher et al., 1991, 2000). Thus, selective feedback perturbation turns an otherwise hidden relational contribution into a diagnostic divergence between independent-item and relation-constrained predictions.

We tested this prediction using selective post-encoding feedback perturbation. After observers encoded two sequentially presented orientations, we provided biased feedback for one orientation and asked whether the resulting bias would remain local to that orientation or would also be expressed in the other orientation, for which feedback had not been provided. Because the two orientations were never seen together and observers reported only individual orientation values, their signed angular relation was neither available in a simultaneous perceptual display nor explicitly required for report. Relation-consistent transfer would therefore indicate that WM had constructed a relation across separately encoded item values and used it to constrain item-value inference.

Using this paradigm, we showed across three experiments that feedback-induced bias transferred to the other orientation in a relation-preserving manner. Experiment 1 established transfer in the direction predicted by the signed angular relation; Experiment 2 showed that transfer was strongest for nearby orientations and attenuated with increasing angular separation, consistent with decreasing relational precision (Chen & Levi, 1996; Howe & Purves, 2005; Lew & Vul, 2015); and Experiment 3 showed that transfer was bidirectional, arguing against a one-way mapping tied to presentation order. Together, these findings suggest that continuous feature values of individual items in visual WM are embedded within a relational structure that constrains how each value is inferred, supporting a view of visual WM as a structured representational system rather than a store of isolated feature estimates.

## Experiment 1

### Method

#### Participants

Participants aged 20–35 years with normal or corrected-to-normal vision were recruited via the online platform Prolific. Before the main experiment, participants completed a brief prescreening session after receiving the task instructions. Participants whose mean absolute error in the prescreening session was 25° or greater were not invited to the main experiment and were compensated £1.50 for their time. One participant was excluded at the prescreening stage.

Each main experiment involved approximately 40–50 min of active task engagement, excluding self-paced breaks between blocks, and participants received £9 for completing it. A total of 24 participants completed Experiment 1. For the main data analysis, one participant was excluded because of extremely low performance, defined as a median absolute error greater than 20° across trials. This criterion was chosen to exclude participants whose performance approached the error expected if they submitted the randomly initialized response orientation without adjustment. Because the response orientation was initialized within ±45° of the target, submitting the initial orientation without adjustment would yield an expected absolute error of 22.5°. As a result, the final analyzed samples included 23 participants (mean age = 29.3 years, 12 females).

The target sample sizes were determined with reference to prior two-orientation continuous-report WM studies. For example, Bae and Luck (2017) used 16 participants in each of three experiments with a closely related paradigm in which observers remembered two sequentially presented orientations and reproduced both after a delay. Because the present feedback-transfer manipulation was novel, no directly relevant prior effect-size estimate was available for a formal a priori power analysis. We therefore aimed to recruit sample sizes comparable to or larger than those used in prior studies with similar two-orientation reproduction procedures. Sensitivity power analyses, conducted for paired-samples tests, indicated that the final analyzed samples provided 80% power at two-tailed *α* = .05 to detect within-subject effects of approximately Cohen’s *d_z_* = .61 in Experiment 1 (Faul et al., 2007; Lakens, 2013). This detectable effect size falls between Cohen’s conventional benchmarks for medium and large effects, suggesting that the final samples were appropriate for detecting medium-to-large within-subject effects in the present design (Cohen, 2013).

All participants provided informed consent online before beginning the experiment. The study was approved by the Institutional Review Board of Seoul National University (IRB No. 2511/003-029) and was conducted in accordance with institutional guidelines for research with human participants.

#### Stimuli

##### display-size calibration

At the beginning of the experiment, participants completed an on-screen size-calibration procedure. They adjusted the size of a gray rectangle until it matched the size of a standard credit card held against the screen. Participants were instructed that they could use any card of the same size, such as a membership card or driver’s license, or use a ruler to set the rectangle width to 85.6 mm. This calibration was used to scale stimulus dimensions in physical units on each participant’s display. Participants were also instructed to maintain an approximate viewing distance of 60 cm throughout the experiment.

##### Orientation targets

Target stimuli consisted of two colored line segments presented sequentially (**Fig. 2A**). Each line measured 5 cm in length and 0.06 cm in width. The first line was orange (RGB: 255, 128, 0), and the second was cyan (RGB: 0, 255, 255). All stimuli were presented at the center of the screen within a light gray circular frame (RGB: 200, 200, 200; diameter = 5.8 cm), which remained visible throughout the trial. A black fixation dot (diameter = 0.3 cm) was continuously displayed at the center of the screen, and participants were instructed to maintain fixation on this dot.

**Figure 2.**
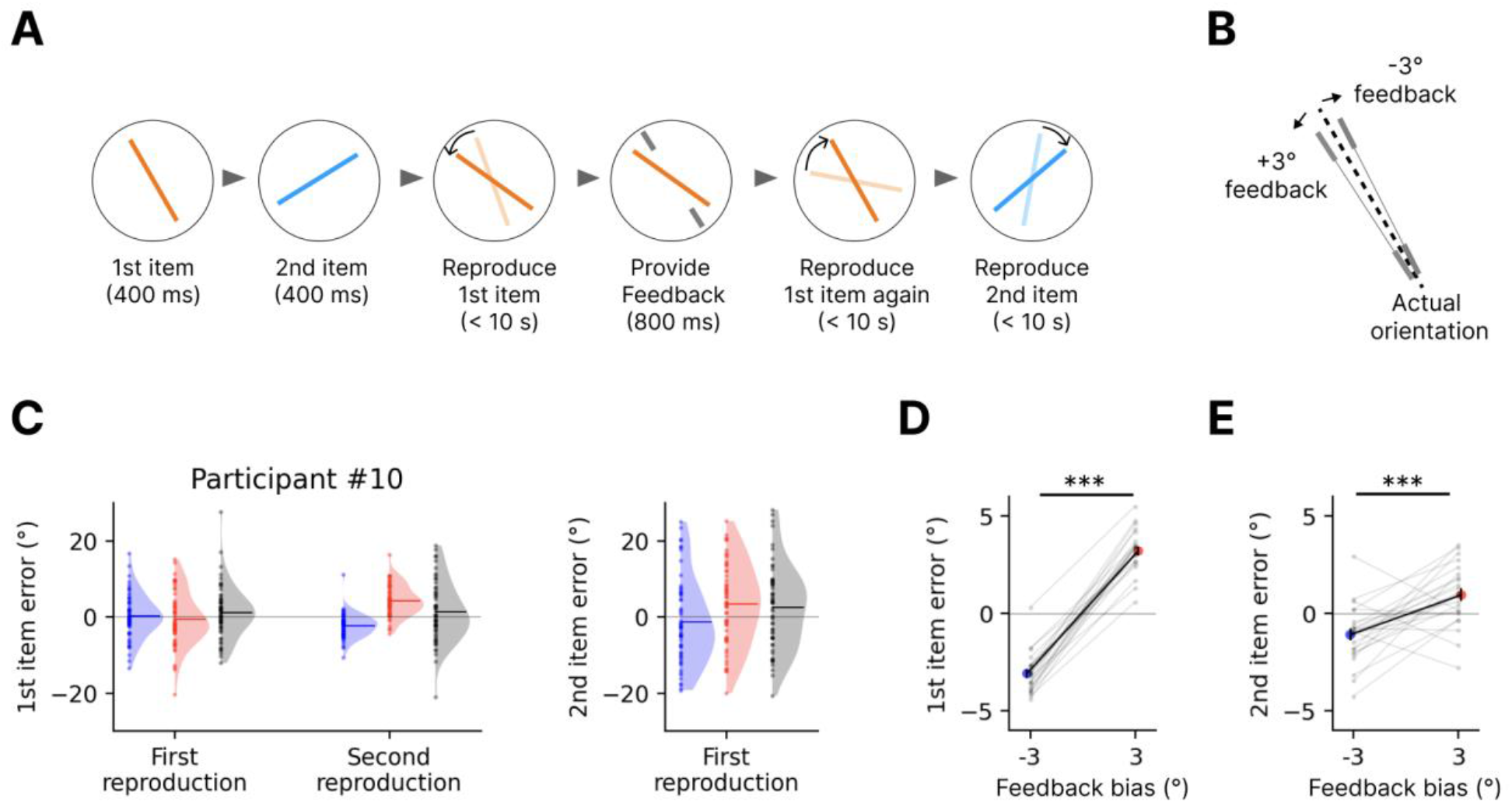
Experimental procedure and feedback-induced transfer in Experiment 1. **(A)** Trial structure. Orange and cyan lines denote the first and second orientations, respectively; response probes were shown in the color of the orientation to be reported, and curved arrows indicate probe adjustment. Participants encoded the two orientations sequentially, reproduced the first orientation, received feedback about it, reproduced it again, and then reproduced the second orientation, for which feedback was not provided. Gray nonius lines denote the feedback display, and gray triangles indicate epoch progression within a trial. **(B)** Feedback manipulation. The black dashed line indicates the actual first orientation, and the gray nonius lines indicate feedback biased by either −3° or +3° relative to that orientation. Only one feedback orientation was presented on each feedback trial. **(C)** Single-trial signed reproduction errors from a representative participant, shown for the first item’s initial and post-feedback reports (left) and the second item’s report (right). Dots indicate individual trials, shaded distributions indicate kernel density estimates, and blue, red, and gray denote the −3°, +3°, and no-feedback conditions, respectively. The thin gray horizontal line indicates zero error. **(D, E)** Group-level signed errors in the second report of the first orientation, for which feedback was provided (**D**), and in the report of the second orientation, for which feedback was not provided (**E**). Gray dots and lines indicate individual participants, colored dots indicate group means, black lines connect the group means, and error bars indicate SEM. ∗, *p* < .05; ∗∗, *p* < .01; ∗∗∗, *p* < .001.

##### Mask

A circular noise mask was used to minimize afterimages. On each trial, a new mask was generated from a 50 × 50 matrix of random grayscale values sampled uniformly from 0 to 255, cropped to a circle and resized to a diameter of 8 cm. A two-dimensional Gaussian filter (SD = 0.06 cm) was applied to smooth the mask.

##### Feedback

It consisted of two short light-gray line segments (RGB: 200, 200, 200) positioned symmetrically around the fixation point. Each segment was 0.5 cm long and 0.06 cm wide, and the two segments were separated by a 4.5 cm gap. The virtual line connecting the segments indicated the feedback orientation.

#### Procedure

Experiment 1 consisted of 10 blocks of 18 trials. Participants could take self-paced breaks between blocks and resume the experiment when ready. Before each block, participants were reminded to maintain an approximate viewing distance of 60 cm from the monitor. At the end of each block, participants were shown a brief performance summary indicating the mean absolute error in the orientation-reproduction task for that block, which was intended to help maintain engagement throughout the experiment.

Each trial began with a brief “Ready” message presented near fixation. The first target line, shown in orange, was presented for 400 ms, followed by a 400 ms interstimulus interval, after which the second target line, shown in cyan, was presented for 400 ms. Participants were instructed to remember both line orientations and their temporal order. After the second line disappeared, a noise mask was presented for 500 ms, followed by an 800 ms blank retention interval.

Participants first made an initial report of the orientation of the first, orange line. A response probe, shown as an orange line matching the color of the to-be-reported target, appeared at an orientation randomly selected within ±45° of the target orientation. The probe color served as an item cue, indicating which of the two remembered orientations should be reported. The ±45° initialization range was chosen to reduce unnecessary adjustment time while still requiring participants to actively adjust the probe on each trial. Participants rotated the probe counter-clockwise or clockwise using the "F" and "J" keys, respectively, and confirmed the adjusted orientation by pressing the "Space bar" key. The response was self-paced, and the probe remained on the screen until the response was confirmed. Participants were instructed to submit their response within 10 s on each trial. If no response was submitted within this time window, the orientation adjusted up to that point was automatically recorded as the trial response.

On feedback trials, which comprised two-thirds of all trials, feedback was presented 200 ms after the initial report. The feedback consisted of two short light-gray line segments indicating the ostensibly correct orientation and was displayed together with the participant’s response line for 800 ms. Participants were instructed that the feedback reflected the true orientation of the first line; in fact, it was systematically biased by either +3° or −3° relative to the actual target orientation (**Fig. 2B**). The three feedback conditions were balanced within each block: each 18-trial block contained six +3° feedback trials, six −3° feedback trials, and six no-feedback trials. On no-feedback trials, feedback segments were not presented, and the response line remained on the screen for the same duration, thereby matching the interval between the initial and second reports across feedback-present and no-feedback trials.

Participants then made a second report of the orientation of the first, orange line, using the same response procedure. This second report was included to assess the immediate effect of the biased feedback on the orientation for which feedback was provided before testing whether the feedback influenced the other item. After this second report, the probe remained visible for 400 ms before disappearing. Participants then made a final report of the orientation of the second, cyan line, using a cyan probe that matched the color of the to-be-reported target. No feedback followed the final report.

The orientation of the first line was randomly sampled on each trial from 18 equally spaced values ranging from 5° to 175° in 10° increments. The orientation of the second line was defined relative to the first-line orientation by adding a signed angular offset randomly sampled from ±10°, ±20°, ±30°, or ±40°.

#### Analysis

The primary dependent variable was signed reproduction error, defined as the angular deviation of the reported orientation from the target orientation. Although errors were analyzed across all three report phases, the critical analysis focused on signed errors in the final report of the second item, for which no feedback was provided. Each trial included three orientation reproduction reports: the first report for the first item, the second report for the first item after feedback, and the report for the second item. Because systematic response drift and lapses could differ across report phases, errors were mean-centered separately for each participant and report phase. Reports with an absolute mean-centered error greater than 30° were excluded only from the corresponding report-phase analysis, rather than causing the entire trial to be rejected. This procedure reduced unnecessary data loss while accounting for phase-specific response bias. These preprocessing steps were applied consistently across all experiments.

The main analysis tested whether a bias introduced to the first item would influence the final report of the second item. If the orientation memories of the two lines were embedded within a relational structure, then a feedback-induced bias in the remembered orientation of the first item should also shift the report of the second item in the corresponding direction. Thus, the +3° and −3° feedback conditions should produce systematically different signed errors not only in the second report of the first item but also in the final report of the second item. In contrast, if feedback remained local to the orientation for which it was provided, the +3° and −3° feedback conditions should differ in the second report of the first orientation, but not in the final report of the second orientation.

To test these predictions, trials were grouped according to feedback direction, either +3° or −3° relative to the actual target orientation of the first item. No-feedback trials were not included in this feedback-direction contrast because they contained no signed feedback bias. For each participant, we first computed the mean signed error in the second report of the first item separately for the +3° and −3° feedback conditions. These values were compared using a paired-samples t test as a manipulation check of whether the biased feedback shifted the second report of the first orientation.

Critically, the same +3° versus −3° comparison was then applied to signed errors in the final report of the second item. A significant difference between the +3° and −3° feedback conditions in this final second-item report would indicate that the feedback-induced bias transferred from the first item to the second item. For descriptive purposes, the feedback-induced bias was quantified for each participant as the difference between mean signed errors in the +3° and −3° feedback conditions, computed separately for the second report of the first item and the final report of the second item.

### Results

Experiment 1 tested whether biased feedback applied to the first item would influence the final report of the second item. To make the core pattern visible at the single-trial level, we first plotted single-trial signed errors, together with their condition-wise distributions, from an example participant (**Fig. 2C**). In the initial report of the first item, errors were broadly distributed around the target orientation, with little separation among trials that would subsequently receive +3° feedback, −3° feedback, or no feedback. In the second report of the first item, a clear condition-dependent pattern emerged: errors in the +3° and −3° feedback conditions shifted toward the corresponding feedback-indicated values and became more tightly clustered around those values, whereas errors in the no-feedback condition remained broadly distributed around the target. This trial-level pattern illustrates that the biased feedback was incorporated as intended into the participant’s subsequent report of the feedback-cued first item. Critically, the same plot also revealed the key transfer pattern in the final report of the second item: although no feedback had been provided for the second item, errors following +3° and −3° feedback to the first item were separated in the corresponding directions. Thus, the single-trial plot provides an intuitive preview of the feedback-transfer effect tested at the group level below.

Having visualized the core pattern in an example participant, we next asked whether the same pattern held across participants. We first verified that the biased feedback was incorporated into the second report of the first orientation. Consistent with the pattern observed in the example participant, group-averaged signed errors in the second report of the first item shifted toward the feedback-indicated values. Mean signed error was 3.19° (SEM = 0.23) in the +3° feedback condition and −3.10° (SEM = 0.22) in the −3° feedback condition. This difference was highly reliable, t(22) = 25.25, p < .001, Cohen’s dz = 5.27, with condition means closely matching the intended feedback biases (**Fig. 2D**).

The critical test was whether this feedback-induced bias was also expressed in the final report of the second item. Again consistent with the example-participant visualization, final second-item reports showed a corresponding separation between the +3° and −3° feedback conditions. Mean signed error was 0.93° (SEM = 0.33) in the +3° feedback condition and −1.11° (SEM = 0.32) in the −3° feedback condition. This difference was reliable, t(22) = 4.34, p < .001, Cohen’s dz = 0.90, indicating that signed errors in the second-item report were systematically separated according to the direction of the feedback applied to the first item, despite the absence of feedback to the second item (**Fig. 2E**). The magnitude of this transfer effect was smaller than the direct feedback effect observed for the first item: the +3° minus −3° difference was 6.29° (SEM = 0.25) for the second report of the first item and 2.04° (SEM = 0.47) for the final report of the second item, t(22) = 9.53, p < .001, Cohen’s dz = 1.99. Thus, the bias introduced to the first item was reliably but only partially reflected in the report of the second item.

To further characterize the source of this second-item effect, we asked whether it contained a feedback-transfer component beyond two expected influences on second-item reports: (i) baseline inter-item bias from the first item and (ii) any stimulus-driven bias from the physical orientation of the feedback display. We estimated the baseline influence of the first item from no-feedback trials and removed this component from second-item errors in feedback trials. We then modeled the residual second-item errors as a function of the feedback orientation relative to the second item, with or without an additional term coding the direction of the feedback bias relative to the first-item target. The model that includes this feedback-bias-direction term provided a better fit than the feedback-orientation-only model, ΔAIC = 17.82, despite the penalty for the additional predictor. Thus, the second-item effect contained a feedback-induced transfer component over and above baseline inter-item bias and stimulus-driven feedback effects (**Supplementary Fig. S1**).

## Experiment 2

In Experiment 2, we asked whether this feedback-induced transfer depends on the angular separation between the two orientations. If the effect arises because the two item memories are linked through a relational structure, then the strength of transfer should depend on the precision with which their signed angular relation is represented. Motivated by prior work suggesting that estimates of angular relations become less precise as angular separation increases (Chen & Levi, 1996; Howe & Purves, 2005; Lew & Vul, 2015), we predicted that feedback-induced transfer would be strongest for nearby orientations and weaker when the two orientations were farther apart. Experiment 2 directly tested this prediction by contrasting a near condition with a pooled far condition.

### Method

#### Participants

Recruitment, prescreening, sample-size rationale, compensation, exclusion criteria, and ethical procedures were the same as in Experiment 1. Two individuals were excluded during prescreening; of the 20 who completed the main experiment, one met the low-performance exclusion criterion, yielding a final sample of 19 participants (mean age = 29.9 years, 11 females). The sample-size target followed the same rationale as in Experiment 1. A sensitivity analysis indicated that the final sample of 19 participants provided 80% power at a two-tailed α of .05 to detect a within-subject effect of dz=0.68.

#### Stimuli

Stimuli and display-size calibration were identical to those in Experiment 1.

#### Procedure

The procedure was identical to that of Experiment 1, except for the manipulation of the orientation difference between the two items. Whereas Experiment 1 sampled relatively small angular distances between the two items (|Δ*θ*| ≤ 40°), Experiment 2 retained ±10° as the near-distance condition and introduced larger offsets of ±50°, ±60°, and ±70°. The ±10° condition was sampled twice as often as each of the larger-distance conditions to provide a stable estimate of transfer at the distance where the effect was expected to be strongest, while allowing us to test whether transfer weakened across a broader range of larger relational distances.

#### Analysis

The primary analysis tested whether feedback-induced transfer to the second item depended on relational distance. Because Experiment 2 was designed to contrast a near-distance condition with a larger-distance regime, trials were grouped into a near condition (±10°) and a far condition (±50°, ±60°, and ±70°). The three larger-separation conditions were combined to provide a stable estimate of transfer at angular separations where the signed angular relation was expected to be represented with lower precision.

As in Experiment 1, the feedback effect was quantified for each participant as the difference between mean signed reproduction errors in the +3° and −3° feedback conditions, calculated as +3° minus −3°. This measure was computed separately for the near and far conditions. The transfer effect was defined as this feedback-direction difference in the final report of the second item. To test whether transfer depended on relational distance, we compared the transfer effect between the near and far conditions using a paired-samples t test.

### Results

Consistent with our prediction, feedback-induced transfer to the second item depended on relational distance (**Fig. 3C**). In the near-distance condition (|Δθ| = 10°), signed errors in the final report of the second item differed reliably between the +3° feedback condition (M = 1.27°, SEM = 0.56) and the −3° feedback condition (M = −0.92°, SEM = 0.49), t(18) = 3.61, p = .002, Cohen’s dz = 0.83. Thus, when the two remembered orientations were nearby, feedback applied to the first item was reliably expressed in the report of the second item. In contrast, in the far-distance condition, which combined larger angular distances (|Δθ| ∈ {50°, 60°, 70°}), signed errors in the second-item report did not differ between the +3° feedback condition (M = −0.10°, SEM = 0.49) and the −3° feedback condition (M = 0.10°, SEM = 0.42), t(18) = −0.29, p = .78, Cohen’s dz = −0.07.

**Figure 3.**
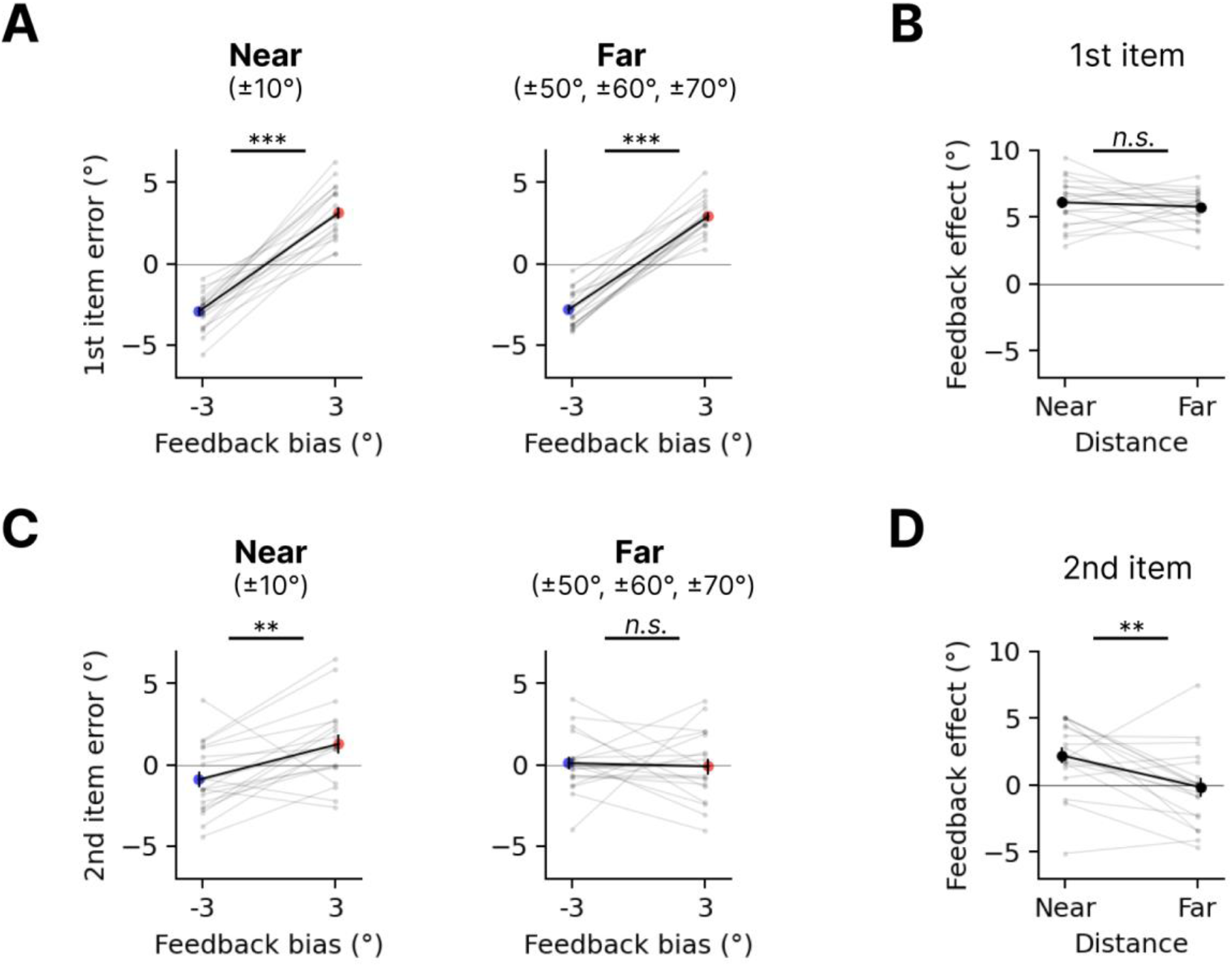
Feedback-induced transfer is attenuated at larger angular separations. **(A)** Signed reproduction errors in the second report of the first orientation, for which feedback was provided, shown separately for near (±10°) and far (±50°, ±60°, and ±70°) angular separations. Blue and red indicate the −3° and +3° feedback conditions, respectively. **(B)** Feedback incorporation effect for the first orientation, quantified as the +3° minus −3° difference in signed reproduction error, shown for the near and far conditions. **(C)** Signed reproduction errors in the report of the second orientation, for which feedback was not provided, shown separately for near and far angular separations. Blue and red indicate the −3° and +3° feedback conditions, respectively. **(D)** Feedback-transfer effect for the second orientation, quantified as the +3° minus −3° difference in signed reproduction error. In all panels, gray dots and lines indicate individual participants. In **A** and **C**, colored dots indicate group means, and black lines connect the group means across feedback conditions. In **B** and **D**, large black dots indicate group means, and black lines connect the near and far conditions. Error bars indicate SEM; thin gray horizontal lines indicate zero error or zero feedback effect; *, p<.05; **, p<.01; ***, p<.001.

A direct comparison confirmed that the transfer effect was larger in the near-distance condition than in the far-distance condition. The feedback-induced separation in second-item errors, quantified as the +3° minus −3° difference, was 2.19° (SEM = 0.61) in the near-distance condition but −0.20° (SEM = 0.69) in the far-distance condition (**Fig. 3D**). The separations differed significantly between the two conditions, t(18) = 3.20, p = .005, Cohen’s dz = 0.73. These results indicate that feedback-induced transfer was robust when the two items were close in orientation space but was markedly attenuated when the relational distance between them was larger.

This distance dependence was not due to weaker incorporation of feedback into the first item at larger distances. The feedback effect for the first item was robust in both the near-distance condition, 6.07°, t(18) = 14.94, p < .001, Cohen’s dz = 3.43, and the far-distance condition, 5.75°, t(18) = 18.98, p < .001, Cohen’s dz = 4.35 (**Fig. 3A**). Moreover, the magnitude of the first-item feedback effect did not differ between the near- and far-distance conditions, t(18) = 0.72, p = .48, Cohen’s dz = 0.16 (**Fig. 3B**). Thus, the reduction in transfer at larger distances cannot be attributed to weaker incorporation of feedback into the first item.

Having established this distance-dependent transfer effect in Experiment 2, we returned to Experiment 1 to ask whether a similar pattern was visible within the narrower range of angular distances used there. Experiment 1 included item pairs separated by 10°, 20°, 30°, and 40°, allowing us to examine transfer across smaller relational distances. This exploratory follow-up analysis showed a qualitatively similar pattern: feedback-induced transfer was evident at smaller angular distances and was weaker toward the largest distance tested in Experiment 1. When these estimates were plotted together with the individual distance conditions from Experiment 2, the combined descriptive pattern suggested a gradual attenuation of transfer as relational distance increased, from robust transfer at nearby orientations to little or no transfer at the larger distances tested in Experiment 2 (**Supplementary Fig. S2**). Although this cross-experiment pattern should be interpreted descriptively, it converges with the primary Experiment 2 result in showing that feedback-induced transfer decreases as angular separation increases, consistent with lower precision of the constructed relation at larger separations.

## Experiment 3

Experiment 3 asked whether this transfer reflects a relational structure linking the two item memories, rather than a fixed one-way mapping anchored to the experienced presentation sequence. In Experiments 1 and 2, feedback was always applied to the first-presented item, which was also the first-reported item. Thus, transfer had so far been tested only in the direction from the first-presented item to the second-presented item.

The design of Experiments 1 and 2—always giving feedback to the first item and probing feedback transfer effects on the second item—leaves open a non-relational account of the observed transfer: the effect may reflect a one-way mapping learned from the experienced presentation sequence rather than a relational structure linking the two remembered values. If WM encodes a forward transformation from the first-presented orientation to the second-presented orientation, feedback about the first orientation could influence the second by applying that experienced mapping. Feedback about the second orientation, however, would require applying the inverse mapping, which was not directly experienced during encoding; transfer should therefore be absent or weaker in the second-to-first direction. By contrast, if the two item values are linked by their signed angular relation within a joint memory state, feedback-induced transfer should occur in both directions relative to presentation order.

Experiment 3 tested this prediction by asking whether feedback-induced transfer is retricted to the experienced first-to-second presentation direction, or whether it also occurs in the reverse, second-to-first direction (**Fig. 4A**).

**Figure 4.**
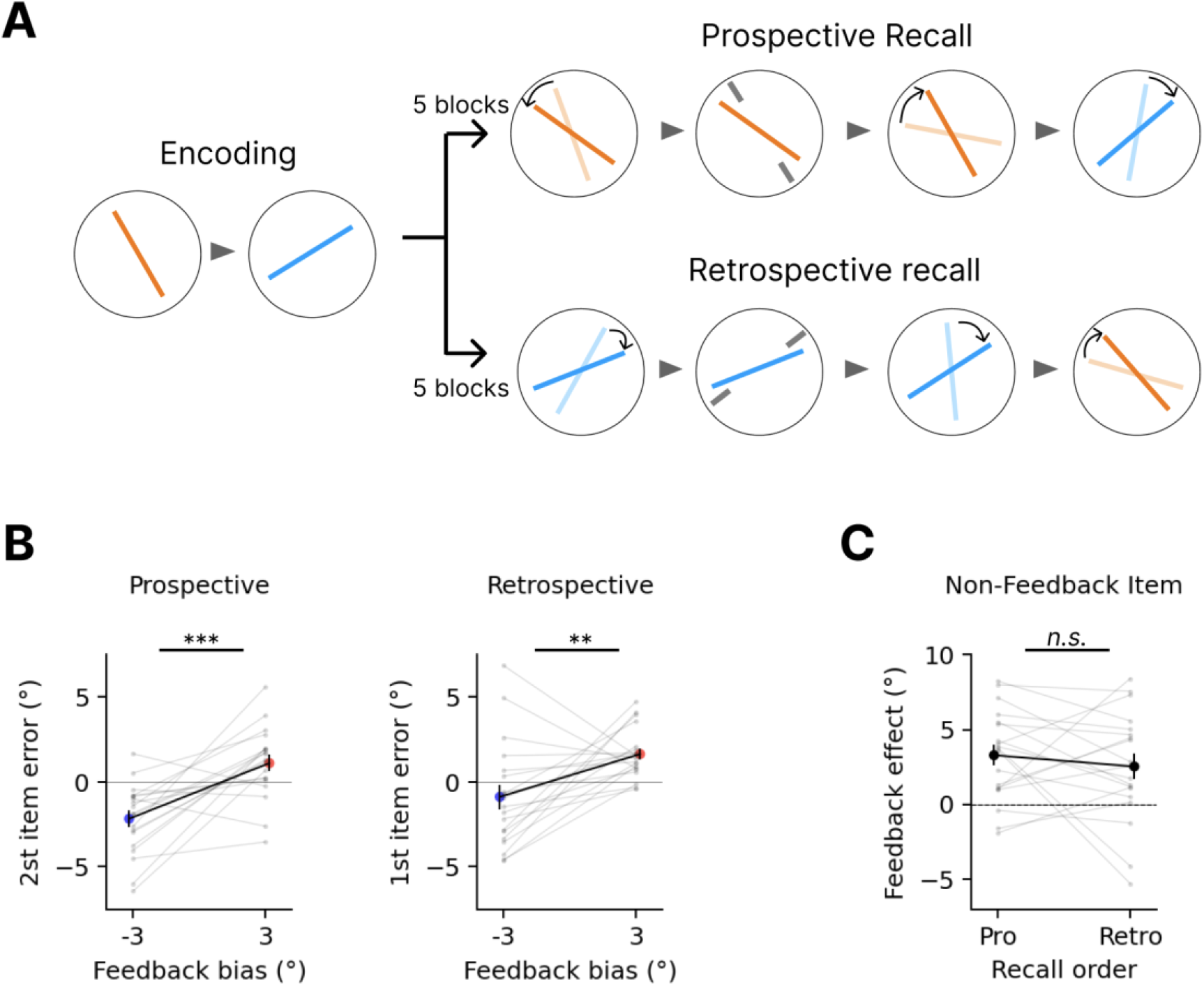
Feedback-induced transfer in the prospective- and retrospective-transfer conditions of Experiment 3. **(A)** Trial structure. Orange and cyan lines denote the first- and second-presented orientations, respectively; response probes were shown in the color of the orientation to be reported, and curved arrows indicate probe adjustment. In the prospective-transfer condition, participants reproduced the first-presented orientation, received feedback about it, reproduced it again, and then reproduced the second-presented orientation. In the retrospective-transfer condition, participants reproduced the second-presented orientation, received feedback about it, reproduced it again, and then reproduced the first-presented orientation. Gray nonius lines denote the feedback display, and gray triangles indicate epoch progression within a trial. **(B)** Signed reproduction errors in the report of the orientation for which feedback was not provided, shown separately for the prospective-transfer condition (left) and retrospective-transfer condition (right). Blue and red indicate the −3° and +3° feedback conditions, respectively. Gray dots and lines indicate individual participants, colored dots indicate group means, and black lines connect the group means across feedback conditions. **(C)** Transfer effects, quantified as the +3° minus −3° difference in signed reproduction error, shown for the prospective- and retrospective-transfer conditions. Gray dots and lines indicate individual participants, large black dots indicate group means, and the black line connects the group means across conditions. Error bars indicate SEM; thin gray horizontal lines indicate zero error or zero transfer effect; *, p<.05; **, p<.01; ***, p<.001.

### Method

#### Participants

Recruitment, prescreening, sample-size rationale, compensation, exclusion criteria, and ethical procedures were the same as in Experiment 1. No individuals were excluded during prescreening; of the 20 who completed the main experiment, one met the low-performance exclusion criterion, yielding a final sample of 19 participants (mean age = 29.0 years, 10 females). The sample-size target followed the same rationale as in Experiment 1. A sensitivity analysis indicated that the final sample of 19 participants provided 80% power at a two-tailed α of .05 to detect a within-subject effect of dz=0.68.

#### Stimuli

Stimuli and display-size calibration were identical to those in Experiment 1.

#### Procedure

The experiment consisted of 10 blocks, divided into two consecutive halves of five blocks each. In the prospective-transfer condition, the order of reporting followed that of presentation: participants reported the first item, received feedback on that item, reported it again, and then reported the second item. This condition therefore tested feedback transfer from the first-presented item to the second-presented item. In the retrospective-transfer condition, the order of reporting was reversed: participants first reported the second-presented item, received feedback on that item, reported it again, and then reported the first-presented item. The order of the prospective- and retrospective-transfer conditions was randomized across participants.

Because the preceding experiments indicated that feedback-induced transfer was most reliably observed at small angular separations, Experiment 3 focused on nearby item pairs to maximize sensitivity to transfer in both directions. The orientation of the second-presented item was defined relative to the first-presented item by a signed offset randomly sampled from ±10°, ±15°, or ±20°. In all other respects, the procedure was identical to that of Experiment 1.

#### Analysis

The primary analysis asked whether feedback-induced transfer was observed not only in the experienced first-to-second presentation direction, but also in the reverse, second-to-first direction. As in the previous experiments, the feedback effect was quantified for each participant as the difference between mean signed reproduction errors in the +3° and −3° feedback conditions, calculated as +3° minus −3°. This measure was computed separately for the prospective-transfer and retrospective-transfer conditions.

In each condition, the transfer effect was defined as the feedback-direction difference in the final report of the item that did not receive feedback. In the prospective-transfer condition, this corresponded to the final report of the second-presented item after feedback had been applied to the first-presented item. In the retrospective-transfer condition, this corresponded to the final report of the first-presented item after feedback had been applied to the second-presented item.

We first tested whether the transfer effect was reliable within each transfer condition. We then compared the magnitude of the transfer effect between the prospective-transfer and retrospective-transfer conditions using a paired-samples t test.

### Results

We first examined whether feedback-induced transfer was present in both the prospective-transfer and retrospective-transfer conditions (**Fig. 4B**). In the prospective-transfer condition, the final report of the second-presented item, which did not receive feedback, showed a reliable separation between feedback conditions. Mean signed reproduction error was 1.11° (SEM = 0.49) in the +3° feedback condition and −2.18° (SEM = 0.48) in the −3° feedback condition, t(18) = 4.73, p < .001, Cohen’s dz = 1.08. Thus, feedback applied to the first-presented item was reliably reflected in the report of the second-presented item, consistent with the transfer effect observed in Experiments 1 and 2.

In the retrospective-transfer condition, the final report of the first-presented item, which did not receive feedback, also showed reliable feedback-induced separation. Mean signed reproduction error was 1.64° (SEM = 0.34) in the +3° feedback condition and −0.89° (SEM = 0.72) in the −3° feedback condition, t(18) = 2.98, p = .008, Cohen’s dz = 0.68. Thus, transfer was also observed when feedback was applied first to the second-presented item, and the effect was tested in the reverse, second-to-first direction.

We next compared the magnitude of the transfer effect between the two transfer conditions (**Fig. 4C**). The transfer effect, quantified as the +3° minus −3° difference in the final report of the item that did not receive feedback, was 3.28° (SEM = 0.69) in the prospective-transfer condition and 2.53° (SEM = 0.85) in the retrospective-transfer condition. This difference was not statistically reliable, t(18) = 0.82, p = .43.

These results indicate that feedback-induced transfer was not limited to the experienced first-to-second presentation direction. Transfer was reliable both when feedback was applied to the first-presented item and expressed in the report of the second-presented item, and when feedback was applied to the second-presented item and expressed in the report of the first-presented item. This bidirectional pattern is consistent with a relational structure linking the two item memories, rather than a fixed one-way mapping anchored to the experienced presentation sequence.

## Discussion

The present study tested whether a relational structure constructed in WM constrains how human observers remember continuous visual feature values of individual items. After two orientations had been presented sequentially—and thus were never seen together—we provided biased feedback for only one of the remembered orientations. As intended, this feedback bias was well incorporated into subsequent reports of that orientation. Critically, the feedback bias was also expressed in reports of the other orientation, for which feedback had never been provided, in a manner that preserved the signed angular relation linking the two remembered orientations. Because the two orientations had never appeared together in a single display, this offset was not visually given; it had to be constructed by actively relating the two remembered orientations within WM feature space. This feedback transfer via relational structure was not a nonspecific spread of feedback. Instead, its strength was regulated by the distance-dependent precision of the constructed angular relation: transfer was reliable for nearby orientations, where the angular offset was represented with greater precision, but was markedly attenuated at larger angular separations, even though feedback incorporation itself remained robust across angular distances. Transfer also occurred in both the experienced first-to-second direction and the reverse direction, arguing against a one-way transformation anchored to the experienced order and supporting a relational structure that was not bound to the sensory sequence in which the items were experienced. Together, these findings suggest that remembered continuous feature values of individual items in visual WM are not maintained merely as independent estimates but are embedded within a relational structure constructed between sequentially encoded items; this structure guides how the remembered value of one item is reshaped when another item’s memory is updated.

### Selective feedback perturbation reveals relation-constrained item memory

Prior studies of WM for reports of continuous feature values have shown that remembered items are often statistically dependent. Reports can be attracted toward or repelled away from other remembered values (Bae & Luck, 2017; Chunharas et al., 2022; Lively et al., 2021; Scotti et al., 2021), errors in reports of multiple items can be correlated (Corbett, 2017; Orhan & Jacobs, 2013; Son et al., 2020), and model-based accounts have proposed that higher-order or relational information can contribute to individual item reports (Ding et al., 2017). These findings are important because they show that item memories are not completely independent. However, not all forms of dependency have the same implications for relational structure in WM.

In the relation-constrained account proposed here, the critical claim is not simply that one item can bias another. Rather, the claim is that individual item values are coupled within a joint memory state constrained by the signed relation between them. This account therefore predicts a relation-preserving tendency: when one remembered value is perturbed, the other value should be reshaped in a way that preserves the signed offset linking the two items.

From this perspective, attraction and repulsion between item estimates demonstrate interactions between remembered feature values, but they do not by themselves establish that the relation between the items is preserved. Both biases can change the distance between item estimates— compressing or exaggerating their separation—rather than maintaining the signed offset that links them. By contrast, positively correlated report errors are more directly compatible with a relation-preserving representation. If the signed relation between two remembered values is preserved, trial-to-trial deviations in one report should tend to be accompanied by deviations in the same direction in the other, producing positively correlated errors. Compared with independent errors of the same magnitude, these shared deviations leave the difference between the two reports relatively stable.

Even so, correlated report errors are not diagnostic of a relational structure constructed in WM because such dependencies can arise from sources that do not require a represented relation between the remembered items. Nearby feature values may be affected in similar ways by broadly tuned perceptual or mnemonic biases, including serial dependence or adaptation-like influences (Fischer & Whitney, 2014; Pascucci et al., 2023). Orientation reports may also be distorted by feature-specific biases that vary smoothly across orientation space, such as cardinal biases, thereby inducing covariance between reports without requiring an underlying relational structure (Girshick et al., 2011; Wei & Stocker, 2015). Correlated errors may also reflect shared variability among feature-selective neural populations, or decision- and response-level history effects during sequential reports, rather than a relational constraint on memory itself (Ryu & Lee, 2017; Smith & Kohn, 2008; H. Zhang & Alais, 2020). Thus, existing report dependencies are compatible with relational structure, but they do not by themselves imply that a constructed relation directly constrains individual item memory.

The methodological advance of the present study lies precisely at this point. Feedback perturbation allowed us to test whether relational structure constrained item memory, rather than merely characterize static dependencies among final reports. Because the perturbation had a known signed direction, its expression in the other remembered orientation provided a diagnostic test of whether the two values were coupled by their signed angular relation. The critical evidence was therefore not merely that the two reports were dependent, but that an experimentally induced change in one orientation was expressed in the other in a relation-preserving manner. Related perturbation approaches have shown that biasing an external reference can shift the estimate of a remembered item (Zamboni et al., 2016); here, selective feedback to one remembered orientation allowed us to test whether such a perturbation would propagate through an internally represented relation to another orientation.

This relation-preserving interpretation also distinguishes the present account from other forms of structured WM, such as ensemble or cluster-based representations (Brady & Alvarez, 2011, 2015; Brady & Tenenbaum, 2013; Corbett, 2017; Orhan & Jacobs, 2013; Son et al., 2020). Different forms of non-independent memory structure should generate different patterns of feedback-induced change. If the two item values were constrained primarily by a pair mean, then a feedback-induced positive update to one item should induce a compensatory negative shift in the other item, thereby preserving the mean. By contrast, if the two item values are coupled within a joint memory state constrained by their signed angular relation, then a positive update to one item should shift the estimate of the other item in the same direction, thereby preserving the offset between them. The observed transfer followed this relation-preserving pattern: reports of the orientation for which feedback had not been provided shifted in the feedback-defined direction, consistent with a relational constraint on the joint memory state rather than a compensatory adjustment to preserve the pair mean.

### Constructing relational structure across sequentially encoded item values

The preceding section characterized the present feedback effect as a relation-preserving tendency: when one remembered orientation was perturbed by feedback, the other orientation shifted in a direction consistent with preserving the signed offset between the two item values. A further implication of the present paradigm concerns how that relation was obtained. Prior studies of configurational and relational memory have shown that visual WM can represent relations among items and use them to support performance, for example when observers detect whether a configuration has been preserved or violated, or when surrounding items provide relational context for a cued target (Bateman et al., 2018; Dent, 2009; Jiang et al., 2000). In many such paradigms, however, the relevant relation is available in the stimulus display, or is reintroduced by the probe display. The present paradigm asked a different question: whether a relation that was not visually given in a simultaneous display could be constructed within WM and then constrain the remembered value of an individual item.

As in previous studies of interactions between sequentially presented orientation values (Bae & Luck, 2017; Ding et al., 2017), the two orientations in our task were also presented sequentially and were never seen together. The signed angular offset linking them was therefore not supplied by visual configuration. Nor was it supplied by a relational probe at test. Instead, the relevant relation had to be constructed by actively relating two separately encoded item values within WM feature space. Sequential presentation alone, however, is not sufficient to demonstrate such construction, because dependencies among final reports can arise from non-relational sources, as discussed above. The critical feature of the present design was the combination of sequential presentation and selective post-encoding perturbation: biased feedback was provided for only one remembered orientation, and the resulting change was measured in the other orientation, for which feedback had not been provided.

This combination of sequential presentation and selective feedback perturbation allowed us to distinguish a visually given relation from a constructed relation that operates within WM. The feedback transfer suggests that the signed angular offset was not merely computed after the two item reports were generated, but was part of the memory structure that constrained how individual item values were estimated. In this sense, the present findings extend prior work on relational and configurational memory by showing that a metric relation between continuous feature values can be constructed across sequentially encoded items and can guide how one item value is reshaped when another, relationally linked item is perturbed.

### A metric, precision-weighted relational constraint

The present findings further characterize the kind of relation that constrained item memory. A relation can be represented categorically or ordinally—for example, which item is larger or which orientation is more clockwise than another (Clevenger & Hummel, 2014; Ding et al., 2017)— without specifying the precise distance between the two values. In the 30° separation condition in Experiment 1 (See **Supplementary Fig. S2**), the feedback-induced shift was not large enough to reverse which orientation was more clockwise. Thus, the ordinal relation would have been preserved even if the orientation for which feedback had not been provided had remained unchanged. Its observed shift therefore cannot be explained by preservation of ordinal order alone. Instead, the relation-preserving direction of transfer is consistent with a constraint imposed by the signed angular offset, which specifies both the direction and the distance linking the two remembered orientations within feature space.

Experiment 2 provided a second signature of this metric constraint. Distance-dependent dependencies have previously been observed in multiple-item reproduction and correlated-error studies (Bae & Luck, 2017; Ding et al., 2017; Orhan & Jacobs, 2013; Son et al., 2020). Here, we used angular distance to test the a priori prediction that feedback transfer would weaken as the constructed relation became less precise over larger separations. Consistent with this prediction, transfer was reliable for nearby orientations but markedly attenuated at larger angular separations. Importantly, incorporation of feedback into the orientation for which it was provided remained robust across distances. The attenuation of transfer therefore cannot readily be attributed to weaker processing or incorporation of feedback for more distant orientation pairs. Nor does it imply that distant orientations were represented without any relation. Rather, it suggests that the constructed relation contributed less strongly to item-value inference as its precision decreased.

This interpretation is consistent with broader evidence that representations of differences, magnitudes, and relational quantities are subject to scale-dependent uncertainty (Lew & Vul, 2015; Sims et al., 2012; Tudusciuc & Nieder, 2010). It is also compatible with empirical-statistical and efficient-coding accounts in which nonuniform environmental statistics shape the precision and bias of visual representations. Natural-scene statistics predict systematic distortions in perceived angular magnitude (Howe & Purves, 2005), and efficient-coding accounts relate orientation biases to inhomogeneity in sensory precision arising from such statistics (Wei & Stocker, 2015). Although the present findings do not identify the source of distance-dependent variation in relational precision, they suggest that relational structure in WM operates not as an all-or-none link between remembered items, but as a metric, precision-weighted constraint whose contribution to item-value inference weakens as angular separation increases.

### Distinguishing relational transfer from non-relational alternatives

Several non-relational accounts could, in principle, produce a shift in reports of the orientation for which feedback had not been provided. We therefore considered whether feedback-induced transfer could be explained by a uniform recalibration of the reporting rule, a nonspecific sensory influence of the oriented feedback display, or a one-way mapping anchored to presentation order.

One possibility is that biased feedback recalibrated an observer’s general reporting rule. Under this account, +3° and −3° feedback would induce corresponding positive and negative shifts in subsequent reports, regardless of which remembered orientation was queried. A uniform, feature-independent recalibration of this kind predicts that the feedback-induced shift should be expressed similarly across the two reported orientations and should not depend selectively on the angular relation between them. Instead, reports of the orientation for which feedback was provided closely incorporated the imposed feedback bias, whereas reports of the other orientation showed only partial transfer, even when that transfer was reliable (**Fig. 2D,E**; **Fig. 3A–D**; **Fig. 4B,C**). Moreover, in Experiment 2, feedback incorporation remained robust across angular distances, whereas transfer was reliable for nearby orientations but markedly attenuated for distant orientations (**Fig. 3A–D**). This dissociation argues against a common shift applied indiscriminately to subsequent reports. It does not exclude every form of history-dependent bias, some of which may themselves vary with feature distance; rather, it rules out the specific account that feedback produced a uniform recalibration of the reporting rule.

Another possibility is that transfer reflected a nonspecific sensory influence of the physical orientation shown in the feedback display. Visual input presented after encoding can bias existing memory representations even when that input was not part of the original memory set (Rademaker et al., 2015; Teng & Kravitz, 2019). Because the feedback display contained an oriented visual signal, reports of the other orientation might have been biased by that physical orientation itself, rather than by the direction of feedback bias defined relative to the orientation for which feedback was provided. This account predicts that, after baseline inter-item bias is taken into account, errors in reports of the other orientation should be explained by the physical feedback orientation. In Experiment 1, however, residual errors after removal of the baseline inter-item bias estimated from no-feedback trials were better fit by a model that included the signed direction of feedback bias than by a model based on the physical feedback orientation alone (ΔAIC = 17.82; **Supplementary Fig. S1**). Thus, the transfer effect cannot be reduced to a nonspecific sensory bias induced by the oriented feedback display.

A third alternative concerns the directionality of transfer. Because Experiments 1 and 2 tested transfer only from the first-presented orientation to the second-presented orientation, the effect could have reflected a fixed one-way mapping established by the experienced presentation sequence rather than a relational structure linking the two remembered values. This account predicts that transfer should be restricted to, or at least substantially stronger in, the experienced first-to-second direction. Experiment 3 did not support this prediction. Feedback-induced transfer was reliable both from the first-presented orientation to the second-presented orientation and in the reverse direction (**Fig. 4B**), and its magnitude did not differ reliably between the prospective and retrospective transfer conditions (**Fig. 4C**). The presence of reliable transfer in both directions argues against a fixed one-way transformation anchored to presentation order.

Taken together, the failure of these alternative accounts refines our claim by specifying what the observed feedback transfer is not: it is not a uniform shift across reports, not driven by the physical feedback orientation itself, and not tied to a fixed first-to-second mapping imposed by the sensory presentation sequence. What remains is a more specific pattern: transfer followed the signed angular relation between the two remembered orientations, weakened with increasing angular distance in a manner consistent with reduced relational precision, and occurred in both directions relative to presentation order. This pattern supports the interpretation that feedback about one remembered orientation influenced the other through the relational structure linking the two item values in WM.

### A probabilistic joint-posterior account of relation-constrained item memory

The preceding sections identified several properties that a successful account of the present findings must explain: feedback-induced change was transferred selectively to another remembered orientation, followed the signed angular relation between the two item values, varied with the precision of that relation, and occurred in both directions relative to presentation order. We next consider how these properties could arise within a probabilistic inference framework. In this framework, observers do not have direct access to the orientations that were presented, but must infer their latent values from uncertain evidence retained in WM. Recent work likewise suggests that WM representations contain distributional information, including uncertainty, beyond a single point estimate (Bays et al., 2024; Li et al., 2021; Zhou et al., 2025). Our aim is not to propose a uniquely identified or fitted mechanism, but to provide a principled probabilistic description of how relational information could constrain individual item values. At the level of the joint memory state, we contrast an independent-item posterior, in which the two item values are uncoupled, with a relation-constrained posterior, in which joint item states are weighted according to their compatibility with the remembered signed relation. One possible generative model and corresponding probabilistic inference scheme through which these posterior structures could arise is developed in **Supplementary Text S3**.

The principal consequence of this difference is visible in the geometry of the joint posterior over the two latent orientations, *p*(*θ*_1_, *θ*_2_ ∣ ***y_M_***) (**Fig. 5**). In the independent-item model, the available evidence contains only item-specific mnemonic evidence, and the joint posterior factorizes:

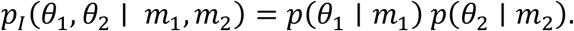

The resulting probability distribution contains no coupling between the two latent item values: Information about one orientation provides no information about the other (**Fig. 5A**). In the relation-constrained model, the same item-specific evidence remains available, but WM additionally constructs and preserves relational information about the signed offset, *θ*_2_ − *θ*_1_. The joint posterior therefore no longer factorizes into separate item-specific posteriors. Instead, this relational constraint introduces posterior dependence between the two item values and elongates the probability distribution along combinations of *θ*_1_and *θ*_2_that preserve the remembered offset (**Fig. 5B**). Thus, in this model, relational information does not merely coexist with item-specific evidence for the two orientations; it changes which combinations of item values are jointly probable.

In a simplified schematic formulation,

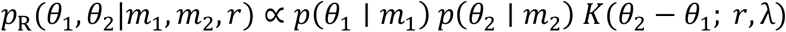

where *r* denotes the mnemonic evidence of the signed angular relation, and *K* is a relation-constraining function of the offset *θ*_2_ − *θ*_1_that weights joint orientation states according to how closely their offset matches *r*, with λ controlling the sharpness of this constraint. This schematic expression implements a simple compatibility-weighting logic for combining item-specific and relational information and characterizes the dependence structure of the joint posterior without specifying the underlying generative and inferential processes through which such a joint posterior could arise. **Supplementary Text S3** develops one plausible generative model and corresponding probabilistic inference scheme through which such a relation-constrained posterior could arise.

**Figure 5.**
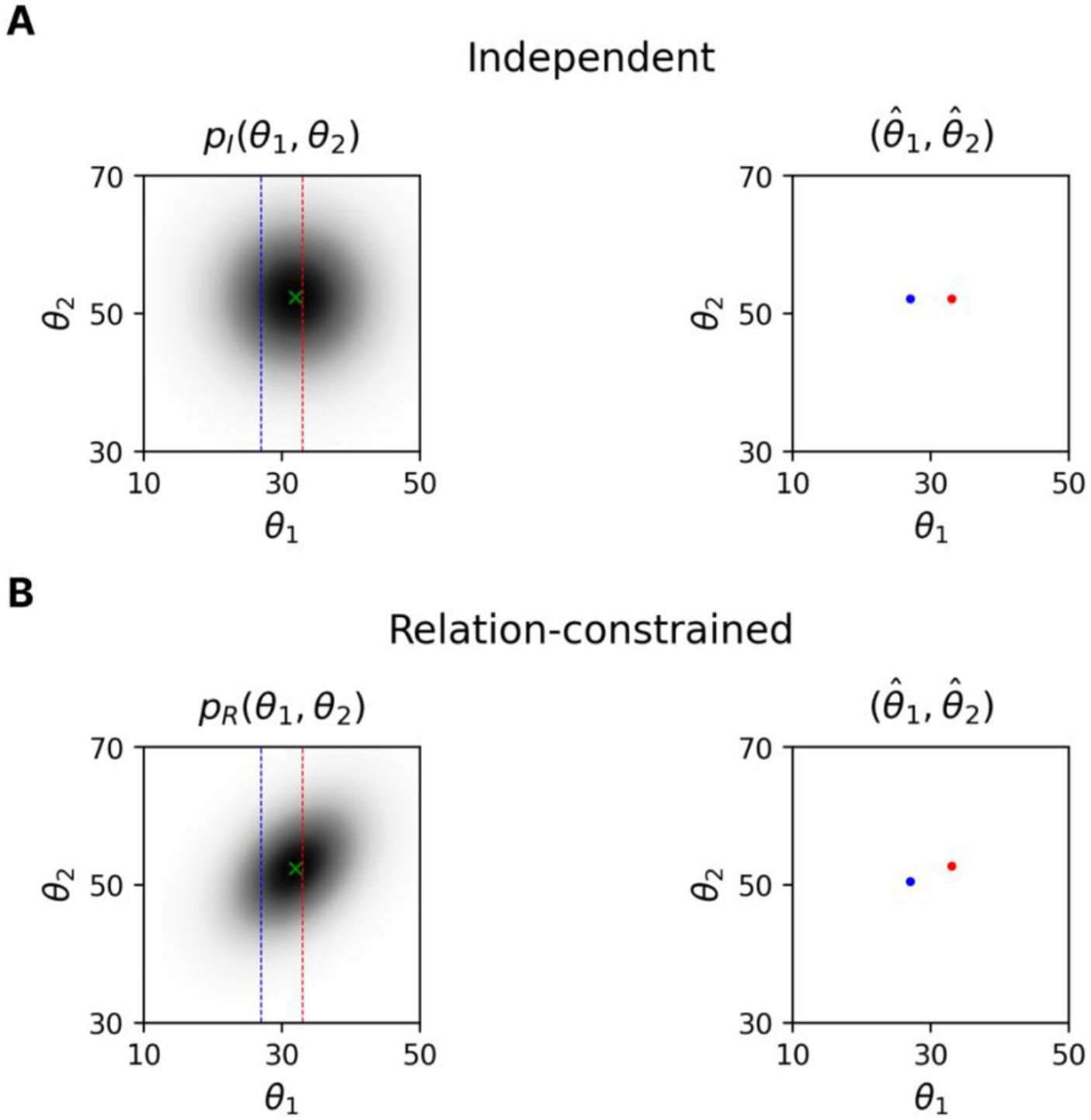
Probabilistic account explaining feedback transfer. **(A)** Independent-item model. The joint posterior factorizes into the item-specific posteriors, *p_I_*(*θ*_1_, *θ*_2_|*m*_1_, *m*_2_) = *p*(*θ*_1_|*m*_1_)*p*(*θ*_2_|*m*_2_), producing an approximately axis-aligned distribution (left). The true values of θ_1_ and θ_2_ are 30^∘^ and 50^∘^, respectively, whereas the center of the remembered posterior, indicated by the green cross, is displaced by sensory and memory noise. Feedback specifying the value of θ_1_ is equivalent to conditioning the joint posterior on that value, or taking a slice of the posterior at the corresponding position along the θ_1_axis. The blue and red dashed lines indicate negatively biased (27^∘^) and positively biased (33^∘^) feedback, respectively. The right panel shows the resulting estimates, 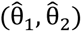, with blue and red dots corresponding to the negative- and positive-feedback conditions. Because the two items are represented independently, feedback changes 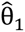 but leaves 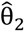 unchanged. **(B)** Relation-constrained model. Plotting conventions are the same as in panel A. Mnemonic evidence about the signed angular relation induces posterior dependence between the two item values, elongating the joint posterior *p_R_*(*θ*_1_, *θ*_2_|*m*_1_, *m*_2_, *r*) along combinations of θ_1_ and θ_2_ that preserve their remembered offset (left). Consequently, conditioning on negatively versus positively biased feedback for θ_1_ produces different estimates of 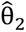 (right). Thus, conditioning on feedback about one orientation shifts the estimate of the other orientation in a relation-preserving direction, producing feedback transfer.

Feedback has different consequences under these two posterior geometries (**Fig. 5**). Feedback is not modeled as overwriting the original mnemonic evidence. Rather, it is treated as specifying a value for one latent orientation, and the already-formed joint posterior is conditioned on the value specified by feedback. In the independent-item model, conditioning on one orientation leaves the posterior estimate of the other unchanged because the two dimensions are uncoupled. In the relation-constrained model, conditioning on one orientation restricts the tilted joint posterior to a different slice and therefore shifts the conditional estimate of the other orientation in a relation-preserving direction. Because the other orientation remains constrained by its own item-specific evidence, this shift need not match the magnitude of the shift in the orientation for which feedback was provided; the same conditioning logic applies regardless of which orientation receives feedback, allowing transfer in both directions.

The relation-constrained joint posterior also extends this account to the graded variation in feedback transfer (**Supplementary Fig. S5**). Because a broader *K* imposes a weaker constraint on the joint posterior, transfer weakens as uncertainty in the remembered relation increases. If this uncertainty increases with angular separation—that is, if *K* becomes broader, or equivalently *λ* decreases, at larger separations—the account would predict the attenuation of transfer with increasing angular separation observed in Experiment 2.

The same relation-constrained posterior geometry that accounts for feedback transfer can also explain earlier findings that relation-preserving changes are relatively difficult to detect (Bateman et al., 2018; Jiang et al., 2000; Orhan & Jacobs, 2013). When item-wise deviations are matched, a relation-preserving change moves the two-item state along the elongated, relatively high-probability ridge of the joint posterior. The changed state therefore remains more probable under the remembered distribution and provides weaker evidence for change. By contrast, a relation-distorting change moves away from this high-probability ridge into a lower-probability region because it violates the remembered relation, thereby providing stronger evidence for change (**Supplementary Fig. S6**).

Taken together, these considerations provide a probabilistic account of the refined set of findings identified above. When the two remembered orientations are represented independently, conditioning on feedback about one orientation leaves the posterior estimate of the other unchanged. When mnemonic evidence about the signed angular relation couples the two values within a joint posterior, however, the same conditioning operation produces a relation-preserving shift in the other orientation. The magnitude of this shift depends on uncertainty in the remembered relation, and the same conditioning logic applies in either direction. Thus, the observed transfer need not require a separate rule for transferring bias between item estimates. Instead, it follows from inference over a joint memory state whose geometry is structured by the signed angular relation linking the two item values.

## Conclusion

Prior work has established that reports of multiple item values in visual WM are often statistically dependent, but such dependencies do not by themselves show that a relation constructed between remembered feature values directly constrains the representation of each value. By selectively perturbing one of two sequentially encoded orientations, the present study provides such evidence. The feedback-induced change in one orientation was expressed in the other in a manner that preserved the signed angular relation linking the two remembered orientations, demonstrating that separately encountered item values can become coupled within a relation-constrained joint memory state. The graded and bidirectional nature of this transfer further indicates that the constructed relation operates not as an all-or-none association or an order-bound transformation, but as a metric constraint on item-value inference whose influence varies with the precision of the remembered relation.

These findings invite a broader view of visual WM—not simply as a store of individual feature estimates, but as a workspace in which relations are actively constructed and used to constrain how remembered values are updated when new information becomes available. An important direction for future research will be to determine when such relational structures are formed, how flexibly they generalize across items and feature domains, and how they interact with item-specific evidence to guide memory-based inference. Addressing these questions may clarify how relational structure supports flexible inference in WM.

## Acknowledgements

This work was supported by Korea Basic Science Institute (National Research Facilities and Equipment Center) grant funded by the Ministry of Education (RS-2024-00435727) and the National Research Foundation of Korea (NRF), funded by the Ministry of Science and ICT (RS-2024-00349515).

## Author contributions

Conceptualization, J.L. and S.-H.L.; Methodology, J.L. and S.-H.L.; Investigation, J.L.; Writing – Original Draft, J.L. and S.-H.L.; Writing – Review & Editing, J.L. and S.-H.L.; Funding Acquisition, S.-H.L.; Supervision, S.-H.L.

## References

Alvarez, G. A., & Oliva, A. (2009). Spatial ensemble statistics are efficient codes that can be represented with reduced attention. Proceedings of the National Academy of Sciences, 106(18), 7345–7350. 10.1073/pnas.0808981106

Ariely, D. (2001). Seeing Sets: Representation by Statistical Properties. Psychological Science, 12(2), 157–162. 10.1111/1467-9280.00327

Bae, G.-Y., & Luck, S. J. (2017). Interactions between visual working memory representations. Attention, Perception, & Psychophysics, 79(8), 2376–2395. 10.3758/s13414-017-1404-8

Bartlett, F. C. (1995). Remembering: A study in experimental and social psychology. Cambridge university press.

Bateman, J. E., Ngiam, W. X., & Birney, D. P. (2018). Relational encoding of objects in working memory: Changes detection performance is better for violations in group relations. Plos One, 13(9), e0203848.

Bays, P. M., & Husain, M. (2008). Dynamic Shifts of Limited Working Memory Resources in Human Vision. Science, 321(5890), 851–854. 10.1126/science.1158023

Bays, P. M., Schneegans, S., Ma, W. J., & Brady, T. F. (2024). Representation and computation in visual working memory. Nature Human Behaviour, 8(6), 1016–1034. 10.1038/s41562-024-01871-2

Behrens, T. E., Muller, T. H., Whittington, J. C., Mark, S., Baram, A. B., Stachenfeld, K. L., & Kurth-Nelson, Z. (2018). What is a cognitive map? Organizing knowledge for flexible behavior. Neuron, 100(2), 490–509.

Brady, T. F., & Alvarez, G. A. (2011). Hierarchical Encoding in Visual Working Memory: Ensemble Statistics Bias Memory for Individual Items. Psychological Science, 22(3), 384–392. 10.1177/0956797610397956

Brady, T. F., & Alvarez, G. A. (2015). Contextual effects in visual working memory reveal hierarchically structured memory representations. Journal of Vision, 15(15), 6. 10.1167/15.15.6

Brady, T. F., & Tenenbaum, J. B. (2013). A probabilistic model of visual working memory: Incorporating higher order regularities into working memory capacity estimates. Psychological Review, 120(1), 85–109. 10.1037/a0030779

Chen, S., & Levi, D. M. (1996). Angle judgment: Is the whole the sum of its parts? Vision Research, 36(12), 1721–1735.

Chong, S. C., & Treisman, A. (2003). Representation of statistical properties. Vision Research, 43(4), 393–404.

Chunharas, C., Rademaker, R. L., Brady, T. F., & Serences, J. T. (2022). An adaptive perspective on visual working memory distortions. Journal of Experimental Psychology: General, 151(10), 2300–2323. 10.1037/xge0001191

Clevenger, P. E., & Hummel, J. E. (2014). Working memory for relations among objects. Attention, Perception, & Psychophysics, 76(7), 1933–1953. 10.3758/s13414-013-0601-3

Cohen, J. (2013). Statistical power analysis for the behavioral sciences. routledge.

Corbett, J. E. (2017). The Whole Warps the Sum of Its Parts: Gestalt-Defined-Group Mean Size Biases Memory for Individual Objects. Psychological Science, 28(1), 12–22. 10.1177/0956797616671524

Dent, K. (2009). Coding categorical and coordinate spatial relations in visual–spatial short-term memory. Quarterly Journal of Experimental Psychology, 62(12), 2372–2387. 10.1080/17470210902853548

D’Esposito, M., & Postle, B. R. (2015). The cognitive neuroscience of working memory. Annual Review of Psychology, 66(1), 115–142.

Ding, S., Cueva, C. J., Tsodyks, M., & Qian, N. (2017). Visual perception as retrospective Bayesian decoding from high-to low-level features. Proceedings of the National Academy of Sciences, 114(43). 10.1073/pnas.1706906114

Faul, F., Erdfelder, E., Lang, A.-G., & Buchner, A. (2007). G* Power 3: A flexible statistical power analysis program for the social, behavioral, and biomedical sciences. Behavior Research Methods, 39(2), 175–191.

Fischer, J., & Whitney, D. (2014). Serial dependence in visual perception. Nature Neuroscience, 17(5), 738–743. 10.1038/nn.3689

Gentner, D. (1983). Structure-mapping: A theoretical framework for analogy. Cognitive Science, 7(2), 155–170.

Girshick, A. R., Landy, M. S., & Simoncelli, E. P. (2011). Cardinal rules: Visual orientation perception reflects knowledge of environmental statistics. Nature Neuroscience, 14(7), 926–932. 10.1038/nn.2831

Haberman, J., & Whitney, D. (2007). Rapid extraction of mean emotion and gender from sets of faces. Current Biology, 17(17), R751–R753.

Halford, G. S., Wilson, W. H., & Phillips, S. (2010). Relational knowledge: The foundation of higher cognition. Trends in Cognitive Sciences, 14(11), 497–505. 10.1016/j.tics.2010.08.005

Hemmer, P., & Steyvers, M. (2009). A Bayesian Account of Reconstructive Memory. Topics in Cognitive Science, 1(1), 189–202. 10.1111/j.1756-8765.2008.01010.x

Howe, C. Q., & Purves, D. (2005). Natural-scene geometry predicts the perception of angles and line orientation. Proceedings of the National Academy of Sciences, 102(4), 1228–1233.

Huttenlocher, J., Hedges, L. V., & Duncan, S. (1991). Categories and particulars: Prototype effects in estimating spatial location. Psychological Review, 98(3), 352.

Huttenlocher, J., Hedges, L. V., & Vevea, J. L. (2000). Why do categories affect stimulus judgment? Journal of Experimental Psychology: General, 129(2), 220.

Jiang, Y., Olson, I. R., & Chun, M. M. (2000). Organization of visual short-term memory. Journal of Experimental Psychology: Learning, Memory, and Cognition, 26(3), 683.

Kemp, C., & Tenenbaum, J. B. (2008). The discovery of structural form. Proceedings of the National Academy of Sciences, 105(31), 10687–10692.

Lakens, D. (2013). Calculating and reporting effect sizes to facilitate cumulative science: A practical primer for t-tests and ANOVAs. Frontiers in Psychology, 4, 62627.

Lew, T. F., & Vul, E. (2015). Ensemble clustering in visual working memory biases location memories and reduces the Weber noise of relative positions. Journal of Vision, 15(4), 10. 10.1167/15.4.10

Li, H.-H., Sprague, T. C., Yoo, A. H., Ma, W. J., & Curtis, C. E. (2021). Joint representation of working memory and uncertainty in human cortex. Neuron, 109(22), 3699–3712.e6. 10.1016/j.neuron.2021.08.022

Lively, Z., Robinson, M. M., & Benjamin, A. S. (2021). Memory Fidelity Reveals Qualitative Changes in Interactions Between Items in Visual Working Memory. Psychological Science, 32(9), 1426–1441. 10.1177/0956797621997367

Luck, S. J., & Vogel, E. K. (1997). The capacity of visual working memory for features and conjunctions. Nature, 390(6657), 279–281. 10.1038/36846

Oberauer, K. (2009). Design for a working memory. Psychology of Learning and Motivation, 51, 45–100.

Orhan, A. E., & Jacobs, R. A. (2013). A probabilistic clustering theory of the organization of visual short-term memory. Psychological Review, 120(2), 297.

Parkes, L., Lund, J., Angelucci, A., Solomon, J. A., & Morgan, M. (2001). Compulsory averaging of crowded orientation signals in human vision. Nature Neuroscience, 4(7), 739–744.

Pascucci, D., Tanrikulu, Ö. D., Ozkirli, A., Houborg, C., Ceylan, G., Zerr, P., Rafiei, M., & Kristjánsson, Á. (2023). Serial dependence in visual perception: A review. Journal of Vision, 23(1), 9. 10.1167/jov.23.1.9

Rademaker, R. L., Bloem, I. M., De Weerd, P., & Sack, A. T. (2015). The impact of interference on short-term memory for visual orientation. Journal of Experimental Psychology: Human Perception and Performance, 41(6), 1650–1665. 10.1037/xhp0000110

Ryu, J., & Lee, S.-H. (2017). Stimulus-Tuned Structure of Correlated fMRI Activity in Human Visual Cortex. Cerebral Cortex, cercor;bhw411v1. 10.1093/cercor/bhw411

Scotti, P. S., Hong, Y., Leber, A. B., & Golomb, J. D. (2021). Visual working memory items drift apart due to active, not passive, maintenance. Journal of Experimental Psychology: General, 150(12), 2506–2524. 10.1037/xge0000890

Sims, C. R., Jacobs, R. A., & Knill, D. C. (2012). An ideal observer analysis of visual working memory. Psychological Review, 119(4), 807–830. 10.1037/a0029856

Smith, M. A., & Kohn, A. (2008). Spatial and Temporal Scales of Neuronal Correlation in Primary Visual Cortex. The Journal of Neuroscience, 28(48), 12591–12603. 10.1523/JNEUROSCI.2929-08.2008

Son, G., Oh, B.-I., Kang, M.-S., & Chong, S. C. (2020). Similarity-based clusters are representational units of visual working memory. Journal of Experimental Psychology: Learning, Memory, and Cognition, 46(1), 46–59. 10.1037/xlm0000722

Summerfield, C., Luyckx, F., & Sheahan, H. (2020). Structure learning and the posterior parietal cortex. Progress in Neurobiology, 184, 101717.

Tenenbaum, J. B., Kemp, C., Griffiths, T. L., & Goodman, N. D. (2011). How to grow a mind: Statistics, structure, and abstraction. Science, 331(6022), 1279–1285.

Teng, C., & Kravitz, D. J. (2019). Visual working memory directly alters perception. Nature Human Behaviour, 3(8), 827–836. 10.1038/s41562-019-0640-4

Tervo, D. G. R., Tenenbaum, J. B., & Gershman, S. J. (2016). Toward the neural implementation of structure learning. Current Opinion in Neurobiology, 37, 99–105.

Tudusciuc, O., & Nieder, A. (2010). Comparison of length judgments and the Müller-Lyer illusion in monkeys and humans. Experimental Brain Research, 207(3–4), 221–231. 10.1007/s00221-010-2452-7

Van den Berg, R., Shin, H., Chou, W.-C., George, R., & Ma, W. J. (2012). Variability in encoding precision accounts for visual short-term memory limitations. Proceedings of the National Academy of Sciences, 109(22), 8780–8785.

Wei, X.-X., & Stocker, A. A. (2015). A Bayesian observer model constrained by efficient coding can explain “anti-Bayesian” percepts. Nature Neuroscience, 18(10), 1509–1517. 10.1038/nn.4105

Wilken, P., & Ma, W. J. (2004). A detection theory account of change detection. Journal of Vision, 4(12), 11. 10.1167/4.12.11

Wyble, B., Tam, J., Deal, I., & Bowman, H. (2025). Understanding the flexibility of working memory: Compositionality, generative processing, anchors and holistic representations. Neuroscience & Biobehavioral Reviews, 179, 106387. 10.1016/j.neubiorev.2025.106387

Zamboni, E., Ledgeway, T., McGraw, P. V., & Schluppeck, D. (2016). Do perceptual biases emerge early or late in visual processing? Decision-biases in motion perception. Proceedings of the Royal Society B: Biological Sciences, 283(1833), 20160263. 10.1098/rspb.2016.0263

Zhang, H., & Alais, D. (2020). Individual difference in serial dependence results from opposite influences of perceptual choices and motor responses. Journal of Vision, 20(8), 2–2.

Zhang, W., & Luck, S. J. (2008). Discrete fixed-resolution representations in visual working memory. Nature, 453(7192), 233–235. 10.1038/nature06860

Zhou, Y., Curtis, C. E., Fougnie, D., & Sreenivasan, K. K. (2025). Probabilistic working memory representations in human cortex guide behavior. bioRxiv, 2025.11. 17.688881.

