## Supplementary material for "Relational Structure Constrains Individual Value Estimates in Visual Working Memory": Supplemtal Texts and Figures

**Supplementary Text S1. Separating feedback-induced transfer from baseline inter-item and feedback-stimulus effects**

The final report of the non-feedback (e.g., second item) could be influenced by multiple factors (**Fig. S1A**). First, even in the absence of feedback, the first item may bias reports of the second item through baseline inter-item interactions. Second, because the feedback display itself contained an oriented visual signal, the physical orientation of the feedback could also exert a nonspecific influence on the subsequent second-item report, either through attraction toward or repulsion away from the feedback orientation. These influences are not mutually exclusive with the feedback-transfer effect tested in Experiment 1. Rather, the key question is whether the direction of the feedback bias applied to the first item explains additional variance in second-item errors after accounting for both the baseline influence of the first item and the physical orientation of the feedback itself.

To address this question, we used the population data from Experiment 1. For clarity, we refer to the feedback-cued item as the first item and the item that did not receive feedback as the second item, consistent with the fixed report structure of Experiment 1.

We first estimated the baseline influence of the first item on second-item errors using no-feedback trials. Specifically, we fitted a generalized additive model (GAM) to the reproduction error of the second item as a function of the relative orientation between the first and second items:

E^(2)^*_i_* = *β*_0_ + *s*(X^(1)^*_i_*) + *ε_i_*	(S1),

where E^(2)^ denotes the reproduction error of the second item, X^(1)^ denotes the orientation of the first item relative to the second item, *β*_0_ is the intercept, s(·) is a smooth function, ε is the error term, and *i* indexes trials from the no-feedback condition. The GAM was fitted using the pyGAM package (LinearGAM with a spline term). The smoothing parameter λ was selected via grid search using the default implementation, which optimizes the generalized cross-validation criterion. This model captured baseline inter-item biases that were present independently of feedback (**Fig. S1B)**.

We then used the fitted GAM to remove the baseline first-item influence from second-item errors in feedback trials. If second-item errors were fully explained by the combination of baseline inter-item influence and the physical orientation of the feedback display, then the residual second-item errors should be accounted for by the feedback orientation relative to the second item. We therefore compared two models of the residual second-item error.

The first model included only the physical feedback orientation relative to the second item:

R^(2)^*_j_* = *β*_0_ + *s*(F*_j_*) + *ε_j_*	(S2),

where R^(2)^ denotes the residual second-item error after removing the baseline influence of the first item estimated from Eq. (S1), F denotes the feedback orientation relative to the second item, and *j* indexes trials from the feedback condition.

The second model included the same feedback-orientation term but additionally included the direction of the feedback bias relative to the first-item target, that is, whether the feedback was biased by +3° or −3° relative to the actual orientation of the first item:

R^(2)^*_j_* = *β*_0_ + *s*(F*_j_*) + *β*_1_C*_j_* + *ε_j_*	(S3),

where C is a categorical variable indicating the direction of the feedback bias relative to the first item.

Eq. (S2) captures the contribution of the physical feedback orientation after accounting for baseline inter-item influence (**Fig. S1C**). Eq. (S3) tests whether second-item errors are additionally explained by how the feedback was applied to the first item (**Fig. S1D**). Thus, the comparison between these models asks whether the second-item effect contains a feedback-induced transfer component beyond the baseline inter-item bias and the nonspecific influence of the feedback stimulus itself.

The model including feedback-bias direction provided a better fit than the feedback-orientation-only model, Eq. S1-2: AIC = 19421.06; Eq. 3: AIC = 19403.24; ΔAIC = 17.81. Thus, second-item errors were not fully explained by baseline inter-item influence together with a nonspecific effect of the physical feedback orientation. Instead, after accounting for these factors, second-item errors still depended on how feedback was applied to the first item. This result supports the interpretation that the second-item effect contains a feedback-induced transfer component over and above ordinary inter-item bias and stimulus-driven feedback effects.

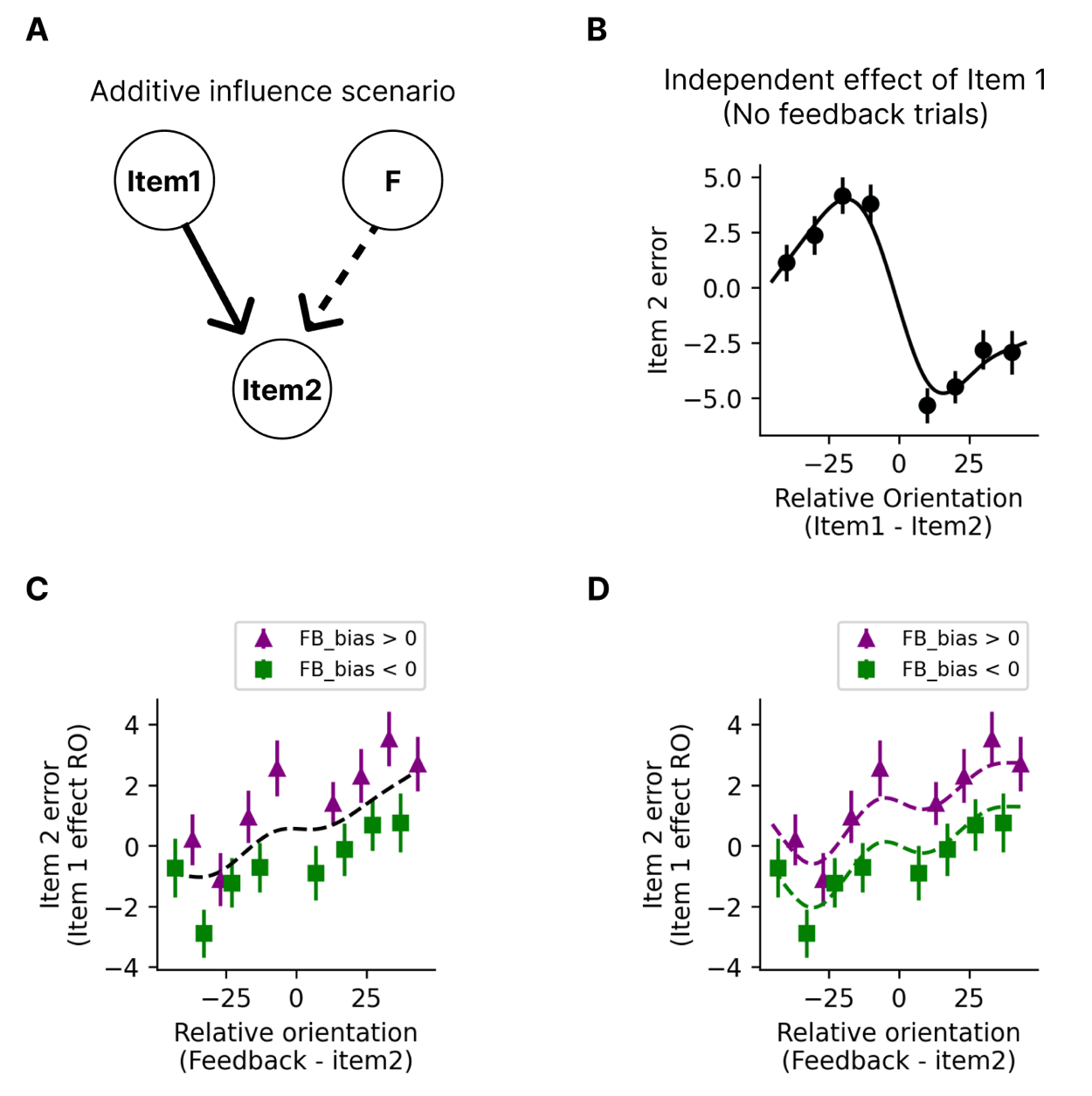

**Figure S1. Separating feedback-induced transfer from baseline inter-item and feedback-stimulus effects.** (A) Schematic of the additive influence scenario. The final report of the second item may be influenced by both the first item and the physical orientation of the feedback display. (B) Baseline influence of the first item on second-item errors, estimated from no-feedback trials using a generalized additive model. (C) Second-item errors in feedback trials plotted as a function of feedback orientation relative to the second item. Dashed line indicates the prediction based on the feedback-orientation-only model after accounting for baseline first-item influence. (D) Residual second-item errors were better captured when the model additionally included feedback-bias direction relative to the first-item target. Error bars indicate ±1 SEM.

**Supplementary Text S2. Distance-dependent attenuation of feedback-induced transfer across Experiments 1 and 2**

The primary analysis of Experiment 2 showed that feedback-induced transfer was stronger for nearby item pairs than for more distant item pairs. To visualize how this transfer varied across individual angular distances, we plotted signed reproduction errors for the item that did not receive feedback separately for the +3° and −3° feedback conditions. This format makes it possible to see directly whether the two feedback conditions produced a separation in the predicted direction at each relational distance.

We first applied this analysis to Experiment 1, which included item pairs separated by 10°, 20°, 30°, and 40°. Although Experiment 1 was not designed as a formal test of distance-dependent transfer, it allowed us to examine whether feedback-induced separation in second-item reports varied across a narrower range of relatively small angular distances. Signed errors in the item that did not receive feedback were more positive following +3° feedback than following −3° feedback at distances of 10°, 20°, and 30°, whereas this separation was no longer reliable at 40°.

We then applied the same visualization to Experiment 2, which contrasted a near-distance condition of 10° with larger-distance conditions of 50°, 60°, and 70°. Consistent with the primary near-versus-far analysis reported in the main text, signed errors in the item that did not receive feedback were clearly separated by feedback-bias direction at 10°, but this separation was not reliable at the larger distances. Thus, when Experiments 1 and 2 were plotted in the same format, the descriptive pattern suggested that feedback-induced transfer was strongest at nearby orientations and weakened as relational distance increased.

This cross-experiment visualization was intended as a descriptive summary rather than a single formal inferential test, because the two experiments differed in their distance ranges and sampling schemes. Nevertheless, the matched format highlights a convergent pattern across experiments: feedback-induced transfer was expressed as a reliable separation between the +3° and −3° feedback conditions at smaller relational distances, and this separation diminished or disappeared at larger distances.

We fitted an ordinary least-squares (OLS) regression model to the demeaned reproduction error of the non-feedbacked item. Because reproduction errors were demeaned within participant, we used an OLS model with participant-clustered standard errors rather than including participant-level random intercepts. The model included corrective-feedback bias condition (−3 vs. +3), absolute difference between the two orientations (10, 20, 30, 40 in Exp 1; 10, 50, 60, 70 in Exp 2), and their interaction as categorical predictors:

*Non-feedbacked item error ~ bias × difference*

Bias condition and difference magnitude were both treated as categorical factors. Standard errors were computed using participant-clustered robust covariance estimates to account for repeated observations within participants. Because the coefficients of this categorical interaction model depend on reference coding, the regression coefficients were not interpreted directly. Instead, we conducted planned simple-effect contrasts comparing the +3 and −3 bias conditions separately at each difference magnitude. P-values from the four planned contrasts were corrected using the Benjamini–Hochberg false discovery rate procedure.

Model-estimated means and standard error of the mean (SEM) are shown in **Fig. S2.**

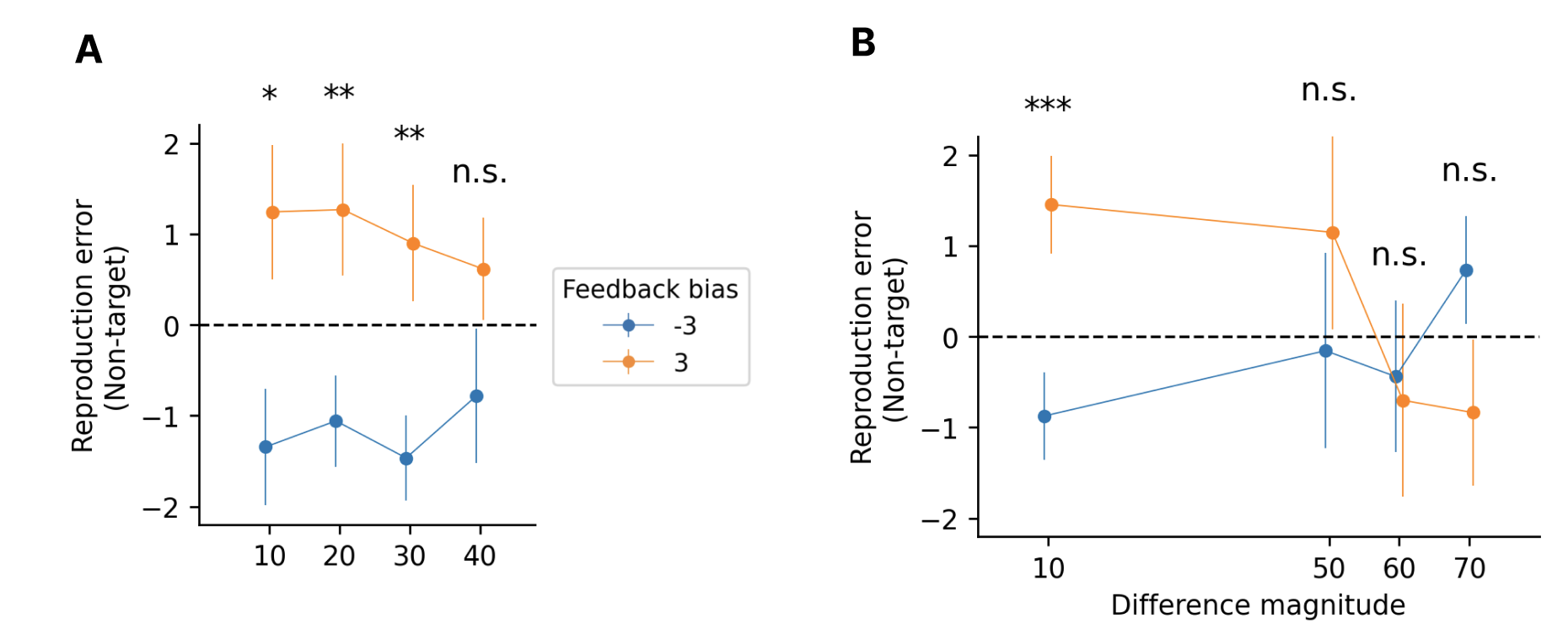

**Figure S2. Distance-dependent attenuation of feedback-induced transfer across Experiments 1 and 2.** **(A)** Signed reproduction errors for the item that did not receive feedback in Experiment 1, plotted as a function of item-pair distance. Experiment 1 included angular distances of 10°, 20°, 30°, and 40°. **(B)** Signed reproduction errors for the item that did not receive feedback in Experiment 2, plotted as a function of item-pair distance. Experiment 2 included a near-distance condition of 10° and larger-distance conditions of 50°, 60°, and 70°. In both panels, errors are plotted separately for trials in which feedback to the other item was biased by +3° or −3° relative to that item’s true orientation. Points indicate group means, and error bars indicate ±1 SEM. Asterisks indicate planned contrasts comparing the +3° and −3° feedback conditions at each distance, with p-values corrected across distances within each experiment using the Benjamini–Hochberg false discovery rate procedure. *pFDR < .05, **pFDR < .01, ***pFDR < .001; n.s., not significant.

#### Supplementary Text S3. A probabilistic inference account of relation-constrained working memory

In the Discussion and **Fig. 5** of the main text, we conceptualize memory for two sequentially presented orientations as a probabilistic joint distribution over possible item states. This Supplement provides one explicit formal route by which such a joint distribution can arise. We first specify evidence-generation models for independent-item and relation-constrained accounts, then derive the joint posterior over the two latent orientations, show how this posterior is conditioned on a feedback-specified value for one orientation and thereby produces transfer to the other orientation, and finally use the same posterior geometry to explain attenuation, distance dependence, potential sources of individual variability, and relation-preserving changes. The goal is not to claim that this is the only possible model, or that the specific assumptions below uniquely determine the observed behavior. Rather, the derivation provides a concrete and internally consistent scenario in which relational evidence reshapes item memory into a coupled joint posterior. Other assumptions about how relational evidence is generated, weighted, or maintained could lead to similar posterior geometries and similar qualitative predictions. The value of the present formulation is that it makes explicit how the qualitative account proposed in the main text can be grounded in a principled probabilistic inference framework.

### S3.1 Evidence generation models for independent-item and relation-constrained accounts

In the orientation reproduction task, the observer does not have direct access to the presented orientations. Instead, the observer has noisy evidence generated from those orientations and must infer the latent world orientations that could have generated that evidence. We contrast two evidence-generation models: an independent-item model and a relation-constrained model.

In both models, the latent world orientations first generate sensory evidence states. These sensory states can be interpreted as noisy sensory measurements generated by the presented orientations. For the two orientations,

$$S_{1}=\theta_{1}+\epsilon_{s1}, S_{2}=\theta_{2}+\epsilon_{s2},$$

where

$$\epsilon_{s1}\mathcal{\sim N}\left( 0,\sigma_{s1}^{2} \right), \epsilon_{s2}\mathcal{\sim N}\left( 0,\sigma_{s2}^{2} \right).$$

The item-specific mnemonic evidence variables are then generated from these sensory evidence states after additional item-specific memory noise:

$$m_{1}=S_{1}+\epsilon_{m1}, m_{2}=S_{2}+\epsilon_{m2},$$

where

$$\epsilon_{m1}\mathcal{\sim N}\left( 0,\sigma_{m1}^{2} \right), \epsilon_{m2}\mathcal{\sim N}\left( 0,\sigma_{m2}^{2} \right).$$

Equivalently,

$$m_{1}=\theta_{1}+\epsilon_{s1}+\epsilon_{m1}, m_{2}=\theta_{2}+\epsilon_{s2}+\epsilon_{m2}.$$

Because $S_{1}$and $S_{2}$ arise from sequentially presented orientations, constructing $\Delta$ from these states does not imply that the relation is visually available in a simultaneous display. Rather, the model assumes that the retained sensory evidence states are related within WM after both have become available. The critical difference between the independent-item and relation-constrained models is whether the observer also constructs mnemonic evidence about the signed angular relation from the sensory evidence states.

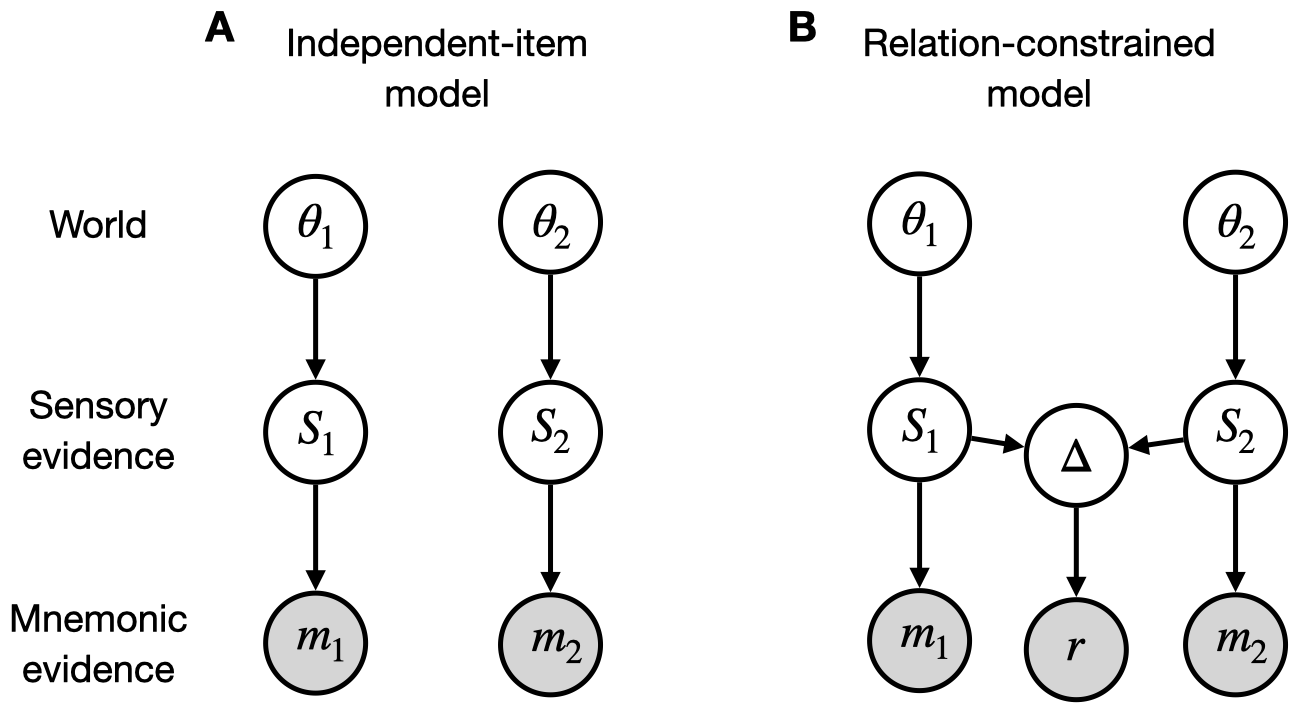

**Figure S3. Generative models for the independent-item and relation-constrained accounts.** **(A)** In the independent-item model, the latent world orientations $\theta_{1}$ and $\theta_{2}$ generate sensory evidence states $S_{1}$ and $S_{2}$, which in turn generate item-specific mnemonic evidence $m_{1}$ and $m_{2}$. No relational variable or relational evidence is constructed. **(B)** In the relation-constrained model, the same item-specific pathways are retained, but the sensory evidence states $S_{1}$ and $S_{2}$ additionally give rise to a relational variable $\Delta$, corresponding to their signed angular relation. This relational variable generates mnemonic evidence $r$ about the signed angular relation. Thus, $r$ is not an independent third observation of the world orientations, nor is it a post hoc subtraction of $m_{2}-m_{1}$. Instead, $r$ is internally constructed mnemonic relational evidence derived from sensory evidence states. Filled nodes indicate observed evidence variables available to the observer, and open nodes indicate latent variables.

#### Independent-item model

In the independent-item model, no relational variable is constructed from $S_{1}$ and $S_{2}$, and no relational evidence is available for inference (**Fig. S3A**). After marginalizing over the sensory evidence states, the available evidence consists only of the two item-specific mnemonic measurements. Let

$$y_{I}=\left[ \begin{matrix} m_{1} \\ m_{2} \end{matrix} \right], \theta=\left[ \begin{matrix} \theta_{1} \\ \theta_{2} \end{matrix} \right].$$

The independent-item observation model can be written as

$$y_{I}=H_{I}\theta+\epsilon_{I},$$

where

$$H_{I}=\left[ \begin{matrix} 1 & 0 \\ 0 & 1 \end{matrix} \right]$$

and

$$\epsilon_{I}=\left[ \begin{matrix} \epsilon_{s1}+\epsilon_{m1} \\ \epsilon_{s2}+\epsilon_{m2} \end{matrix} \right].$$

Thus,

$$\mathbb{E}\left[ y_{I}\mid\theta\right]=H_{I}\theta=\left[ \begin{matrix} \theta_{1} \\ \theta_{2} \end{matrix} \right].$$

Assuming that the sensory and item-specific mnemonic noise terms are mutually independent, the covariance of the independent-item evidence vector is diagonal:

$$\Sigma_{I}=\left[ \begin{matrix} \sigma_{s1}^{2}+\sigma_{m1}^{2} & 0 \\ 0 & \sigma_{s2}^{2}+\sigma_{m2}^{2} \end{matrix} \right].$$

This diagonal covariance expresses the key assumption of the independent-item model: the two item pathways provide separable evidence for the two latent orientations. A relation between the orientations may be computed from the resulting item-specific estimates. However, because it is not represented as mnemonic evidence in this model, it does not constrain inference about either latent orientation.

#### Relation-constrained model

In the relation-constrained model, the same item-specific evidence pathways are retained, but the observer additionally constructs relational evidence from the sensory evidence states (**Fig. S3B**). Throughout this Supplement, the signed relation is defined in the second-minus-first direction. Thus, the relational variable is

$$\Delta=S_{2}-S_{1}.$$

Relational mnemonic evidence $r$ is then generated from this relational variable:

$$r=\Delta+\epsilon_{r},$$

where

$$\epsilon_{r}\mathcal{\sim N}\left( 0,\sigma_{r}^{2} \right).$$

Substituting the sensory evidence states into the relational observation equation gives

$$r=\theta_{2}-\theta_{1}+\epsilon_{s2}-\epsilon_{s1}+\epsilon_{r}.$$

This formulation is intended to capture the idea that relational evidence is constructed from sensory evidence states, not directly and independently from the latent world orientations. The relational evidence $r$ therefore reflects the signed offset from the first orientation to the second orientation, but it is not an independent third measurement of $\theta_{1}$ and $\theta_{2}$. Nor is it a simple subtraction of the item-specific mnemonic evidence $m_{2}-m_{1}$. Rather, $r$ is mnemonic evidence about the signed angular relation $\Delta$, constructed from $S_{1}$ and $S_{2}$.

Thus, in the relation-constrained model, the observation vector is

$$y_{R}=\left[ \begin{matrix} m_{1} \\ m_{2} \\ r \end{matrix} \right].$$

The corresponding observation model is

$$y_{R}=H_{R}\theta+\epsilon_{R},$$

where

$$H_{R}=\left[ \begin{matrix} 1 & 0 \\ 0 & 1 \\ -1 & 1 \end{matrix} \right]$$

and

$$\epsilon_{R}=\left[ \begin{matrix} \epsilon_{s1}+\epsilon_{m1} \\ \epsilon_{s2}+\epsilon_{m2} \\ -\epsilon_{s1}+\epsilon_{s2}+\epsilon_{r} \end{matrix} \right].$$

Therefore,

$$\mathbb{E}\left[ y_{R}\mid\theta\right]=H_{R}\theta=\left[ \begin{matrix} \theta_{1} \\ \theta_{2} \\ \theta_{2}-\theta_{1} \end{matrix} \right].$$

A key consequence of this evidence-generation model is that the observation noise covariance is not diagonal. Because $r$ is generated from $\Delta$, which is constructed from $S_{1}$ and $S_{2}$, it shares sensory-stage noise with the item-specific mnemonic evidence. The covariance of the relation-constrained evidence vector is

$$\Sigma_{R}=\left[ \begin{matrix} \sigma_{s1}^{2}+\sigma_{m1}^{2} & 0 & -\sigma_{s1}^{2} \\ 0 & \sigma_{s2}^{2}+\sigma_{m2}^{2} & \sigma_{s2}^{2} \\ -\sigma_{s1}^{2} & \sigma_{s2}^{2} & \sigma_{s1}^{2}+\sigma_{s2}^{2}+\sigma_{r}^{2} \end{matrix} \right].$$

In particular,

$$\mathrm{Cov}\left( m_{1},r \right)=-\sigma_{s1}^{2}, \mathrm{Cov}\left( m_{2},r \right)=\sigma_{s2}^{2},$$

whereas

$$\mathrm{Cov}\left( m_{1},m_{2} \right)=0$$

when the sensory and mnemonic noise terms are independent between items.

This formulation captures the central difference between the two models in a common observation-model format. In the independent-item model, the available evidence is the two-dimensional vector $y_{I}=\left[ m_{1},m_{2} \right]^{T}$, with diagonal covariance. In the relation-constrained model, the available evidence is augmented to $y_{R}=\left[ m_{1},m_{2},r \right]^{T}$, and its covariance is structured because relational evidence shares sensory-stage uncertainty with item-specific evidence. Thus, relation-constrained memory is formalized as a joint probabilistic evidence state in which item-specific evidence and relational evidence are coupled through their shared sensory-stage uncertainty.

All orientation variables are understood within the relevant circular orientation, with relational differences represented as signed angular differences. The Gaussian formulation provides a local approximation for analytic clarity.

### S3.2 Probabilistic inference from available evidence

Having specified the evidence-generation models, we next describe how the observer uses the available evidence to infer the latent item values. The central object in this section is the joint posterior over the two latent orientations, not the feedback effect itself. The no-feedback estimates are then a natural consequence of this posterior: under a squared-error, or L2, loss function, the optimal report is the posterior mean.

For a given model $M\in\{I,R\}$, let $\mathbf{y}_{M}$ denote the evidence vector available under that model. From the observation models defined above,

$$\mathbf{y}_{M}=H_{M}\boldsymbol{\theta}+\boldsymbol{\varepsilon}_{M},$$

where

$$\boldsymbol{\theta}=\left[ \begin{matrix} \theta_{1} \\ \theta_{2} \end{matrix} \right], \boldsymbol{\varepsilon}_{M}\mathcal{\sim N}\left( \mathbf{0},\Sigma_{M} \right).$$

Thus,

$$p\left( \mathbf{y}_{M}\mid\boldsymbol{\theta} \right)\mathcal{=N}\left( \mathbf{y}_{M};H_{M}\boldsymbol{\theta},\Sigma_{M} \right).$$

Once $\mathbf{y}_{M}$ has been observed and thus held fixed, this same conditional density, regarded as a function of $\boldsymbol{\theta}$, defines the likelihood. Assuming a uniform prior over $\theta_{1}$ and $\theta_{2}$, the posterior is proportional to this likelihood:

$$p\left( \boldsymbol{\theta}\mid\mathbf{y}_{M} \right)\propto exp\left[ -\frac{1}{2}\left( \mathbf{y}_{M}-H_{M}\boldsymbol{\theta} \right)^{T}\Sigma_{M}^{-1}\left( \mathbf{y}_{M}-H_{M}\boldsymbol{\theta} \right) \right].$$

Because the exponent is a quadratic function of $\boldsymbol{\theta}$, completing the square expresses it in the standard form of a multivariate Gaussian and makes the posterior center and covariance explicit. The posterior is therefore Gaussian,

$$p\left( \boldsymbol{\theta}\mid\mathbf{y}_{M} \right)\mathcal{=N}\left( \boldsymbol{\theta;}\boldsymbol{\mu}_{M|y},\Sigma_{M\mid y} \right),$$

with covariance

$$\Sigma_{M\mid y}=\left( H_{M}^{T}\Sigma_{M}^{-1}H_{M} \right)^{-1}$$

and mean

$$\boldsymbol{\mu}_{M|y}=\Sigma_{M\mid y}H_{M}^{T}\Sigma_{M}^{-1}\mathbf{y}_{M}.$$

We denote this posterior mean and covariance as

$$\boldsymbol{\mu}_{M|y}=\left[ \begin{matrix} \mu_{1,M} \\ \mu_{2,M} \end{matrix} \right], \Sigma_{M\mid y}=\left[ \begin{matrix} V_{1,M} & C_{M} \\ C_{M} & V_{2,M} \end{matrix} \right].$$

This notation will be useful in the next section, where feedback is modeled as conditioning this already-formed joint posterior on the value specified by feedback for one latent orientation.

#### Independent-item model

For the independent-item model, applying the general result above to the evidence-generation model specified in **S3.1** above shows that, because $H_{I}=I_{2}$and $\Sigma_{I}$ is diagonal, the posterior factorizes into two item-specific terms:

$$p\left( \theta_{1},\theta_{2}\mid\mathbf{y}_{I} \right)=p\left( \theta_{1}\mid m_{1} \right)p\left( \theta_{2}\mid m_{2} \right).$$

Equivalently,

$$\boldsymbol{\mu}_{I|y}=\left[ \begin{matrix} m_{1} \\ m_{2} \end{matrix} \right]$$

and

$$\boldsymbol{\Sigma}_{I\mid y}=\left[ \begin{matrix} \sigma_{s1}^{2}+\sigma_{m1}^{2} & 0 \\ 0 & \sigma_{s2}^{2}+\sigma_{m2}^{2} \end{matrix} \right].$$

Thus, $C_{I}=0$. The independent-item posterior contains no dependence between the two latent item values. Each item estimate is determined by its own mnemonic evidence, and a relation may be computed from the resulting item-specific estimates, but because it is not represented as mnemonic evidence in this model, it does not constrain inference about either latent orientation.

#### Relation-constrained model

For the relation-constrained model, applying the general result above to the observation model specified in S3.1 gives the joint posterior

$$p\left( \boldsymbol{\theta}|\mathbf{y}_{R} \right)\mathcal{=N}\left( \boldsymbol{\theta};\boldsymbol{\mu}_{R|y},\Sigma_{R\mid y} \right),$$

where

$$\Sigma_{R\mid y}=\left( H_{R}^{T}\Sigma_{R}^{-1}H_{R} \right)^{-1}$$

and

$$\boldsymbol{\mu}_{R|y}=\Sigma_{R\mid y}H_{R}^{T}\Sigma_{R}^{-1}\mathbf{y}_{R}.$$

Unlike the independent-item posterior, this posterior generally does not factorize into two item-specific posteriors. The third row of $H_{R}$ maps the latent item values onto their expected signed difference $\theta_{2}-\theta_{1}$, and the non-diagonal covariance $\Sigma_{R}$ reflects the fact that $r$ shares sensory-stage uncertainty with $m_{1}$ and $m_{2}$. Thus, inference about each orientation is constrained not only by its own item-specific evidence, but also by the other item’s evidence together with the relational evidence linking the two orientations.

The intuition becomes clearer in the symmetric case,

$$\sigma_{s1}^{2}=\sigma_{s2}^{2}=\sigma_{s}^{2}, \sigma_{m1}^{2}=\sigma_{m2}^{2}=\sigma_{m}^{2}.$$

Under this symmetry, posterior uncertainty can be decomposed into a common direction, in which the two orientations vary together and their signed relation is preserved, and a difference direction, in which their signed relation changes. The posterior variances along these two directions are denoted by $V_{\mathrm{common}}$ and $V_{\mathrm{difference}}$, respectively. Relational evidence $r$ provides no information about a common shift of the two orientations, so

$$V_{\mathrm{common}}=\sigma_{s}^{2}+\sigma_{m}^{2}.$$

Along the difference direction, the item-based difference and relational evidence provide two measurements of the same signed relation. Their shared sensory-stage uncertainty remains, whereas their independent mnemonic and relation-specific uncertainties combine by precision weighting, yielding

$$V_{\mathrm{difference}}=\sigma_{s}^{2}+\frac{\sigma_{m}^{2}\sigma_{r}^{2}}{2\sigma_{m}^{2}+\sigma_{r}^{2}}.$$

Transforming back to the individual-item coordinates gives

$$V=\frac{P_{\mathrm{common}}+P_{\mathrm{difference}}}{2}, C=\frac{P_{\mathrm{common}}-P_{\mathrm{difference}}}{2}.$$

Thus, $V$is the average of the posterior variances along the common and difference directions. The covariance $C$is half the excess uncertainty along the common direction over that along the difference direction: common-direction fluctuations move the two orientations together and contribute positively to their covariance, whereas difference-direction fluctuations move them in opposite directions and contribute negatively. Because relational evidence more tightly constrains the difference direction, $V_{\mathrm{difference}}<V_{\mathrm{common}}$, yielding $C>0$.

Accordingly, the posterior covariance takes the form

$$\Sigma_{R\mid y}=\left[ \begin{matrix} V & C \\ C & V \end{matrix} \right],$$

where

$$V=\frac{2\sigma_{s}^{2}\sigma_{m}^{2}+\sigma_{s}^{2}\sigma_{r}^{2}+\sigma_{m}^{4}+\sigma_{m}^{2}\sigma_{r}^{2}}{2\sigma_{m}^{2}+\sigma_{r}^{2}}$$

and

$$C=\frac{\sigma_{m}^{4}}{2\sigma_{m}^{2}+\sigma_{r}^{2}}.$$

Because $r$ provides evidence about the signed relation $\theta_{2}-\theta_{1}$, each orientation can be estimated either directly from its own mnemonic evidence or indirectly from the other item after translating it through the represented relation: $m_{2}-r$ predicts $\theta_{1}$, whereas $m_{1}+r$predicts $\theta_{2}$. In the symmetric case, the direct and relation-based estimates share the same sensory-stage uncertainty; after this shared component is factored out, their independent variances are $\sigma_{m}^{2}$and $\sigma_{m}^{2}+\sigma_{r}^{2}$, respectively, so precision weighting gives the relation-based estimate the weight $w=\sigma_{m}^{2}/\left( 2\sigma_{m}^{2}+\sigma_{r}^{2} \right)$.

The posterior means simplify to

$$\mu_{1,R|y}=\left( 1-w \right)m_{1}+w\left( m_{2}-r \right)$$

and

$$\mu_{2,R|y}=\left( 1-w \right)m_{2}+w\left( m_{1}+r \right),$$

where

$$w=\frac{\sigma_{m}^{2}}{2\sigma_{m}^{2}+\sigma_{r}^{2}}.$$

These expressions show that the posterior mean for $\theta_{1}$ is a compromise between its own mnemonic evidence $m_{1}$ and the relation-based prediction $m_{2}-r$. Similarly, the posterior mean for $\theta_{2}$ is a compromise between its own mnemonic evidence $m_{2}$ and the relation-based prediction $m_{1}+r$. If relational evidence is highly uncertain, $\sigma_{r}^{2}$ is large and $w$ approaches zero, so the posterior means reduce to the independent-item estimates. If relational evidence is more precise, the posterior means are more strongly constrained by the relation.

In this symmetric flat-prior case, the sensory-evidence variance $\sigma_{s}^{2}$ does not appear explicitly in the posterior mean because the sensory-stage noise shared between item evidence and relational evidence cancels algebraically in the mean estimate. This cancellation should not be interpreted as meaning that sensory-stage uncertainty is irrelevant. The sensory-evidence variance remains part of the posterior covariance through $V$, and therefore contributes to the uncertainty of the inferred item values.

The relation-constrained posterior also provides a natural way to distinguish direct item-evidence independence from posterior dependence. Even when $m_{1}$ and $m_{2}$ have no direct covariance in the evidence-generation process, the inferred item values can become coupled because both are inferred from a joint evidence state that includes relational evidence. In the symmetric case above, the off-diagonal posterior covariance is $C>0$; moreover, across repeated realizations of the evidence, the two posterior-mean estimates can covary positively. Thus, item-specific evidence can be independent at the level of direct item pathways, yet posterior beliefs about the two latent item values can be coupled through relation-constrained inference.

So far, we have described how probabilistic inference is performed under each evidence-generation model to obtain the joint posterior over the latent item values and its posterior mean, $\boldsymbol{\mu}_{M|y}$, from the available evidence. In no-feedback trials, or in a standard two-item reproduction task without feedback, this posterior mean provides the memory estimates under an L2 loss function. In feedback trials, the same joint posterior becomes the object that is conditioned on the value specified by feedback for one latent orientation, as described next.

##### S3.3 Feedback as joint posterior conditioning

We next specify how the joint posterior formed from the pre-feedback evidence is conditioned on the value specified by feedback. In feedback trials, the observer is shown a value for one of the remembered orientations. For feedback to the first orientation, we denote the value specified by feedback by $f_{1}$ and condition on

$$\theta_{1}=f_{1}.$$

For feedback to the second orientation, we analogously condition on

$$\theta_{2}=f_{2}.$$

Here, $f_{1}$ and $f_{2}$ denote the values specified by feedback. They are not mnemonic evidence variables generated by the pre-feedback memory state, and feedback is not modeled as replacing $m_{1}$ or $m_{2}$ with corrected mnemonic measurements. The original evidence vector remains the evidence generated by the pre-feedback memory state. Instead, feedback enters the model by fixing one latent orientation to the feedback-specified value within the already formed joint posterior. Whether this conditioning affects inference about the other orientation depends on whether the pre-feedback posterior contains dependence between the two latent item values.

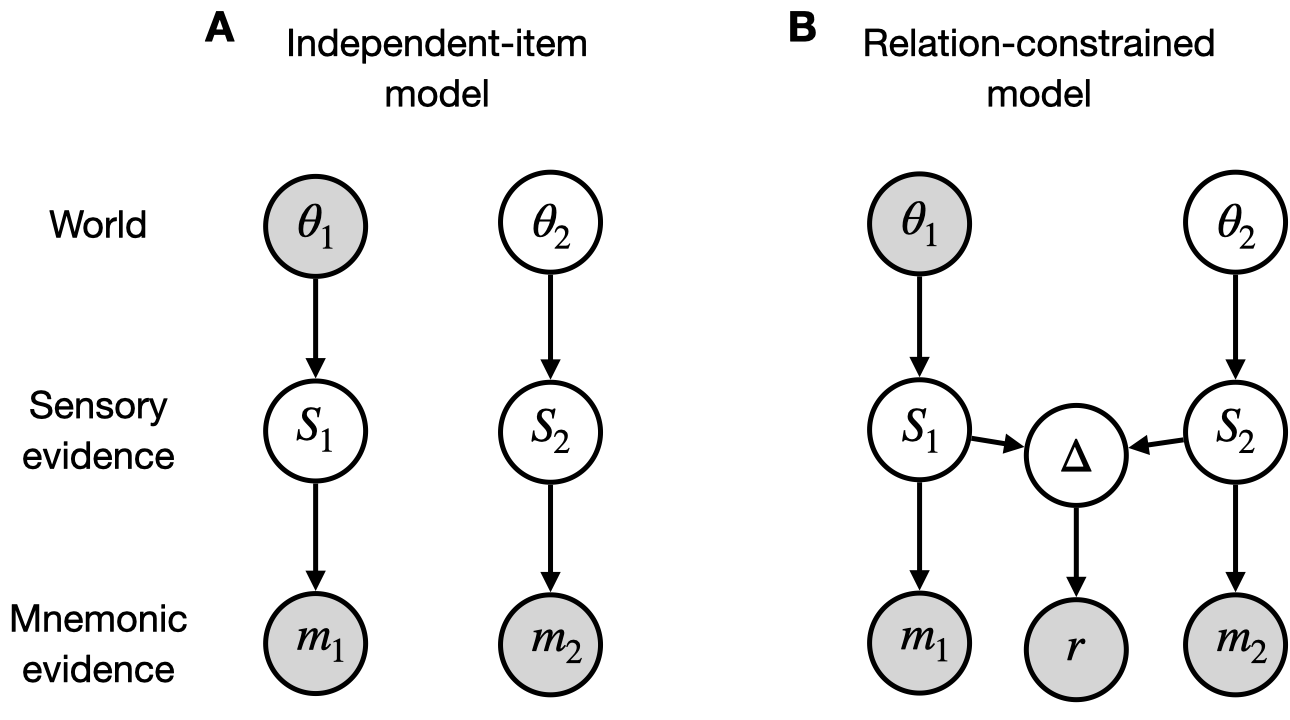

**Figure S4.** **Feedback as conditioning of joint posterior conditioning under the independent-item and relation-constrained models.** **(A)** In the independent-item model, feedback specifies a value $f_{1}$ for the first orientation, the joint posterior is conditioned on $\theta_{1}=f_{1}$. The original item-specific mnemonic evidence variables $m_{1}$ and $m_{2}$ remain available, but because the joint posterior factorizes across items, conditioning on $\theta_{1}=f_{1}$ does not change inference about $\theta_{2}$. **(B)** In the relation-constrained model, the same item-specific evidence variables remain available, and relational evidence $r$ also contributes to the joint posterior over $\theta_{1}$ and $\theta_{2}$. Feedback again specifies a value $f_{1}$for the first orientation, and the joint posterior is conditioned on $\theta_{1}=f_{1}$. Because the posterior over the two item values is coupled, this conditioning changes posterior inference about $\theta_{2}$. Shaded evidence nodes indicate observed evidence variables, whereas the shaded latent node indicates the latent orientation fixed to the feedback-specified value through conditioning. Open nodes indicate unobserved variables. The figure illustrates feedback to the first orientation; feedback to the second orientation is analogous.

**Post-feedback inference about the other orientation with no feedback: a direct derivation from the generative model**

The conditioning illustrated in **Fig. S4** can first be derived directly from the generative model under the feedback constrained. When feedback specifies a value $f_{1}$ for the first orientation, the model treats this value as exact and restricts inference to the conditional state $\theta_{1}=f_{1}$. Inference about the second orientation is then performed within this conditional state, while the observed realization of the original evidence vector $y_{M}$ is held fixed.

In ordinary Bayes’ rule,

$$p\left( A\mid B \right)=\frac{p\left( B\mid A \right)p\left( A \right)}{p\left( B \right)},$$

the posterior plausibility of $A$ after observing $B$ is proportional to two terms: how well $A$ explains the observed evidence, $p\left( B\mid A \right)$, and how plausible $A$ was before observing that evidence, $p\left( A \right)$. The denominator $p\left( B \right)$ normalizes these relative plausibilities into a valid probability distribution. If another condition $C$ is imposed, the same logic is applied within the conditional state defined by $C$. Thus, each term in Bayes’ rule is conditioned on $C$:

$$p\left( A\mid B,C \right)=\frac{p\left( B\mid A,C \right)p\left( A\mid C \right)}{p\left( B\mid C \right)}.$$

In the feedback setting, the corresponding variables are

$$A=\theta_{2}, B=y_{M}, C=\left( \theta_{1}=f_{1} \right).$$

Therefore, applying Bayes’ rule within the feedback-constrained state gives

$$p_{M}\left( \theta_{2}\mid y_{M},\theta_{1}=f_{1} \right)=\frac{p_{M}\left( y_{M}\mid\theta_{2},\theta_{1}=f_{1} \right)p_{M}\left( \theta_{2}\mid\theta_{1}=f_{1} \right)}{p_{M}\left( y_{M}\mid\theta_{1}=f_{1} \right)}.$$

This expression says that, once the constraint $\theta_{1}=f_{1}$ has been imposed, each possible value of $\theta_{2}$ is evaluated according to two factors. The first factor—the first term of the numerator—is how well the observed pre-feedback evidence vector $y_{M}$ would be explained if the latent orientations were $\theta_{1}=f_{1}$ and that particular value of $\theta_{2}$. The second factor—the second term of the numerator—is the prior plausibility of that value of $\theta_{2}$ conditional on $\theta_{1}=f_{1}$but beforeincorporating $y_{M}$. Because the assumed prior factorizes across the two latent orientations,

$$p_{M}\left( \theta_{2}\mid\theta_{1}=f_{1} \right)=p_{M}\left( \theta_{2} \right).$$

The denominator is the normalization constant obtained by integrating the numerator over all possible values of the second orientation:

$$p_{M}\left( y_{M}\mid\theta_{1}=f_{1} \right)=\int p_{M}\left( y_{M}\mid\theta_{2}',\theta_{1}=f_{1} \right)p_{M}\left( \theta_{2}' \right) d\theta_{2}'.$$

Thus, the feedback-constrained posterior can be written as

$$p_{M}\left( \theta_{2}\mid y_{M},\theta_{1}=f_{1} \right)=\frac{p_{M}\left( y_{M}\mid\theta_{1}=f_{1},\theta_{2} \right)p_{M}\left( \theta_{2} \right)}{\int p_{M}\left( y_{M}\mid\theta_{1}=f_{1},\theta_{2}' \right)p_{M}\left( \theta_{2}' \right) d\theta_{2}'}.$$

This is the direct mathematical expression of the conditioning illustrated in **Fig. S4**: the model fixes the first latent orientation at the feedback-specified value $f_{1}$, while the observed pre-feedback evidence vector $\mathbf{y}_{M}$ remains the same and is used to infer the other orientation. As shown in the next subsection, this direct derivation from the generative model is mathematically equivalent to conditioning the already-formed joint posterior on $\theta_{1}=f_{1}$.

**Post-feedback inference about the other orientation with no feedback: an equivalent joint-posterior derivation**

The same conditioning can also be expressed in terms of the joint posterior already formed from the available—pre-feedback—evidence in Section S3.2. We spell out this equivalence because it is the bridge between the graphical interpretation in **Fig. S4** and the feedback-transfer equation below. It does not constitute an additional inference step. Rather, imposing the constraint $\theta_{1}=f_{1}$ directly in the generative formulation and conditioning the already-formed joint posterior on $\theta_{1}=f_{1}$ are two mathematically equivalent descriptions of the same inference.

Before feedback is applied, the available evidence has already produced the joint posterior

$$p_{M}\left( \theta\mid y_{M} \right)=p_{M}\left( \theta_{1},\theta_{2}\mid\mathbf{y}_{M} \right)\mathcal{=N}\left( \theta;\mu_{M\mid y},\Sigma_{M\mid y} \right),$$

where

$$\mu_{M\mid y}=\left[ \begin{matrix} \mu_{1,M} \\ \mu_{2,M} \end{matrix} \right], \Sigma_{M\mid y}=\left[ \begin{matrix} V_{1,M} & C_{M} \\ C_{M} & V_{2,M} \end{matrix} \right].$$

To condition this posterior on the feedback-specified value $f_{1}$, we evaluate its joint density along the slice $\theta_{1}=f_{1}$ and normalize that slice over $\theta_{2}$:

$$p_{M}\left( \theta_{2}\mid\mathbf{y}_{M},\theta_{1}=f_{1} \right)=\frac{p_{M}\left( \theta_{1}=f_{1},\theta_{2}\mid\mathbf{y}_{M} \right)}{p_{M}\left( \theta_{1}=f_{1}\mid\mathbf{y}_{M} \right)}.$$

Using Bayes’ rule, the numerator can be written as

$$p_{M}\left( \theta_{1}=f_{1},\theta_{2}\mid\mathbf{y}_{M} \right)=\frac{p_{M}\left( \mathbf{y}_{M}\mid\theta_{1}=f_{1},\theta_{2} \right)p_{M}\left( \theta_{1}=f_{1},\theta_{2} \right)}{p_{M}\left( \mathbf{y}_{M} \right)}.$$

Under the factorized prior assumed above,

$$p_{M}\left( \theta_{1}=f_{1},\theta_{2} \right)=p_{M}\left( \theta_{1}=f_{1} \right) p_{M}\left( \theta_{2} \right).$$

For fixed $\mathbf{y}_{M}$ and $f_{1}$, the factors $p_{M}\left( \theta_{1}=f_{1} \right)$ and $p_{M}\left( \mathbf{y}_{M} \right)$ do not vary with $\theta_{2}$. The joint-posterior slice is therefore proportional, as a function of $\theta_{2}$, to

$$p_{M}\left( \theta_{1}=f_{1},\theta_{2}\mid\mathbf{y}_{M} \right)\propto p_{M}\left( \mathbf{y}_{M}\mid\theta_{1}=f_{1},\theta_{2} \right)p_{M}\left( \theta_{2} \right).$$

Normalizing this expression over $\theta_{2}$ gives

$$p_{M}\left( \theta_{2}\mid\mathbf{y}_{M},\theta_{1}=f_{1} \right)=\frac{p_{M}\left( \mathbf{y}_{M}\mid\theta_{1}=f_{1},\theta_{2} \right)p_{M}\left( \theta_{2} \right)}{\int_{\Theta} p_{M}\left( \mathbf{y}_{M}\mid\theta_{1}=f_{1},\theta_{2}' \right)p_{M}\left( \theta_{2}' \right) d\theta_{2}'},$$

which is exactly the direct feedback-constrained Bayes expression derived in the preceding subsection. Thus, imposing $\theta_{1}=f_{1}$ in the generative formulation and conditioning the already-formed joint posterior on $\theta_{1}=f_{1}$ yield the same conditional posterior over $\theta_{2}$. Because the orientations are continuous variables, expressions such as $p_{M}\left( \theta_{1}=f_{1}\mid\mathbf{y}_{M} \right)$ denote probability densities evaluated at $f_{1}$, rather than probability masses.

Because the joint-posterior formulation makes the feedback-transfer effect especially transparent and directly uses the posterior quantities derived in Section S3.2, we use this formulation below. Under the Gaussian assumptions, the conditional posterior is also Gaussian. For feedback to the first orientation,

$$\theta_{2}\mid\mathbf{y}_{M},\theta_{1}=f_{1}\mathcal{\sim N}\left( \mu_{2,M}+\frac{C_{M}}{V_{1,M}}\left( f_{1}-\mu_{1,M} \right), V_{2,M}-\frac{C_{M}^{2}}{V_{1,M}} \right).$$

Under a squared-error loss function, the estimate of the other orientation is therefore the conditional posterior mean:

$$\hat{\theta}_{M,2}^{\mathrm{FB}\left( 1 \right)}=\mu_{2,M}+\frac{C_{M}}{V_{1,M}}\left( f_{1}-\mu_{1,M} \right).$$

For feedback to the second orientation, the corresponding estimate is

$$\hat{\theta}_{M,1}^{\mathrm{FB}\left( 2 \right)}=\mu_{1,M}+\frac{C_{M}}{V_{2,M}}\left( f_{2}-\mu_{2,M} \right).$$

These expressions show that feedback transfer is determined jointly by the discrepancy between the feedback-specified value and the pre-feedback posterior mean of the orientation for which feedback was provided, and by the posterior dependence between the two orientations, quantified by $C_{M}/V_{1,M}$ or $C_{M}/V_{2,M}$. If $C_{M}=0$, conditioning on one orientation leaves the estimate of the other unchanged. If $C_{M}>0$, the estimate of the other orientation shifts in the same direction as the feedback discrepancy; if $C_{M}<0$, it shifts in the opposite direction.

###### **Feedback effect in the independent-item model**

Applying the conditional-mean result above to the independent-item posterior, for which $C_{I}=0$, gives, for feedback to the first orientation,

$$\hat{\theta}_{I,2}^{\mathrm{FB}\left( 1 \right)}=\mu_{I,2}.$$

For feedback to the second orientation, the corresponding result is

$$\hat{\theta}_{I,1}^{\mathrm{FB}\left( 2 \right)}=\mu_{I,1}.$$

Thus, conditioning on the feedback-specified value of one orientation leaves the posterior mean of the other orientation unchanged. The model treats the orientation for which feedback was provided as fixed at the specified value, but because the pre-feedback posterior factorizes across the two orientations, this constraint does not propagate to the other orientation.

###### **Feedback effect in the relation-constrained model**

Applying the same conditional-mean result to the relation-constrained posterior, for which $C_{R}$is generally nonzero, gives, for feedback to the first orientation,

$$\hat{\theta}_{R,2}^{\mathrm{FB}\left( 1 \right)}=\mu_{R,2}+\beta_{2\leftarrow1}\left( f_{1}-\mu_{R,1} \right),$$

where

$$\beta_{2\leftarrow1}=\frac{C_{R}}{V_{1,R}}.$$

The gain $\beta_{2\leftarrow1}$specifies how strongly the discrepancy between the feedback-specified value $f_{1}$and the pre-feedback posterior mean $\mu_{1,R}$is expressed in the posterior mean of the other orientation.

For feedback to the second orientation, the corresponding expression is:

$$\hat{\theta}_{R,1}^{\mathrm{FB}\left( 2 \right)}=\mu_{R,1}+\beta_{1\leftarrow2}\left( f_{2}-\mu_{R,2} \right),$$

where

$$\beta_{1\leftarrow2}=\frac{C_{R}}{V_{2,R}}.$$

Thus, feedback transfer depends jointly on the feedback discrepancy and the posterior dependence between the two orientations. In the symmetric case derived above, $C_{R}>0$, so the estimate of the other orientation shifts in the same direction as the feedback discrepancy

The transfer gain can be written explicitly **under the symmetric assumptions used in Section S3.2**:

$$\sigma_{s1}=\sigma_{s2}=\sigma_{s}, \sigma_{m1}=\sigma_{m2}=\sigma_{m}.$$

**Because the two marginal posterior variances are then equal,** $V_{1,R}=V_{2,R}=V$**, and the posterior covariance is** $C_{R}=C$**, the transfer gains in the two directions are identical:**

$$\beta_{2\leftarrow1}=\beta_{1\leftarrow2}\equiv\beta=\frac{C}{V}.$$

**Substituting the expressions for** $\boldsymbol{V}$**and** $\boldsymbol{C}$ **derived in Section S3.2 gives**

$$\beta=\frac{\sigma_{m}^{4}}{\sigma_{m}^{4}+2\sigma_{s}^{2}\sigma_{m}^{2}+\sigma_{r}^{2}\left( \sigma_{s}^{2}+\sigma_{m}^{2} \right)}.$$

The gain $\beta$ is the fraction of the discrepancy between the feedback-specified value and the pre-feedback posterior mean of one orientation that is expressed in the posterior mean of the other orientation. Because the denominator contains the numerator plus additional nonnegative uncertainty terms, $0\leq\beta\leq1$, and $\beta<1$whenever sensory-stage or relation-specific uncertainty is nonzero. Thus, the model naturally predicts attenuated rather than complete feedback transfer.

##### S3.4 Posterior-conditioning gain accounts for attenuated and distance-dependent feedback transfer and identifies potential sources of variability

Section S3.3 showed that feedback transfer in the relation-constrained model is governed by the posterior-conditioning gain. In the symmetric case, the gain is

$$\beta=\frac{\sigma_{m}^{4}}{\sigma_{m}^{4}+2\sigma_{s}^{2}\sigma_{m}^{2}+\sigma_{r}^{2}\left( \sigma_{s}^{2}+\sigma_{m}^{2} \right)}.$$

This expression organizes two principal features of the feedback-transfer effects observed in the experiments—attenuation and dependence on inter-item distance—and identifies potential sources of variability in transfer strength.

First, the gain explains why feedback transfer is attenuated. The denominator equals the numerator plus the nonnegative terms

$$2\sigma_{s}^{2}\sigma_{m}^{2}+\sigma_{r}^{2}\left( \sigma_{s}^{2}+\sigma_{m}^{2} \right).$$

Thus, $0\leq\beta\leq1$, with $\beta<1$whenever sensory-stage or relation-specific uncertainty contributes to inference. The discrepancy between the feedback-specified value and the pre-feedback posterior mean of one orientation is therefore only partially expressed in the posterior mean of the other orientation. Feedback transfer is consequently graded rather than an all-or-none copying of the feedback value.

The feedback values in the experiments were displaced by $3^{\circ}$in opposite directions relative to the presented orientation. Under the approximation that the pre-feedback posterior mean is centered on the presented orientation and that the gain is symmetric across the two feedback directions, the predicted positive-minus-negative feedback contrast for the other orientation is approximately

$$2\times3^{\circ}\times\beta=6^{\circ}\beta.$$

Complete transfer would therefore correspond to an approximately $6^{\circ}$contrast, whereas a contrast of approximately $2^{\circ}$, similar in scale to that observed in the experiments, corresponds to $\beta\approx1/3$.

Second, the same gain provides an account of the distance-dependent transfer observed in Experiment 2. Let $d$ denote the experimentally manipulated absolute angular separation between the two orientations. If relation-specific uncertainty increases with this separation,

$$\sigma_{r}^{2}=g\left( d \right),$$

where $g\left( \cdot\right)$ is an increasing function, then $\beta$ decreases as $d$ increases. Under this assumption, more widely separated orientations are linked by less precise relational evidence, so conditioning on the feedback-specified value of one orientation places a weaker constraint on the other orientation. Distance-dependent transfer therefore follows from the same posterior-conditioning mechanism that produces attenuation.

The gain also shows that different sources of uncertainty have distinct effects on transfer. Increasing relation-specific uncertainty $\sigma_{r}$ decreases transfer by weakening the evidence linking the two orientation values. Increasing sensory-stage uncertainty $\sigma_{s}$ likewise decreases transfer: in the symmetric solution, it increases marginal posterior uncertainty without increasing the covariance that carries the feedback constraint, thereby reducing $C/V$. By contrast, increasing item-specific mnemonic uncertainty $\sigma_{m}$ increases the gain within this parameterization. When an orientation is less strongly anchored by its own mnemonic evidence, the relation-based prediction constrained by feedback has greater influence on its posterior mean.

These dependencies identify potential sources of variability in feedback transfer. Experimental conditions or observers may differ in sensory-stage, mnemonic, or relation-specific uncertainty, producing different posterior-conditioning gains. This is a model implication rather than a result established by the present individual-difference analyses; future work could estimate these uncertainty parameters hierarchically and test the predicted associations directly.

**Fig. S5** illustrates these analytic predictions through parameter sweeps of the posterior-conditioning gain. The curves and landscapes were not fit to the behavioral data. Rather, they show the qualitative regime in which transfer decreases with relation-specific and sensory-stage uncertainty but increases with item-specific mnemonic uncertainty. The strongest transfer is predicted when sensory-stage and relation-specific uncertainty are low while item-specific mnemonic uncertainty is high: the relation remains reliable, whereas the orientation for which feedback was not provided is only weakly anchored by its own mnemonic evidence.

Taken together, the posterior-conditioning gain provides a unified account of the principal qualitative effects. Attenuation follows from $\beta<1$; distance dependence follows if relation-specific uncertainty increases with inter-item angular separation; and variability in sensory-stage, mnemonic, and relational precision provides testable potential sources of variation in transfer strength.

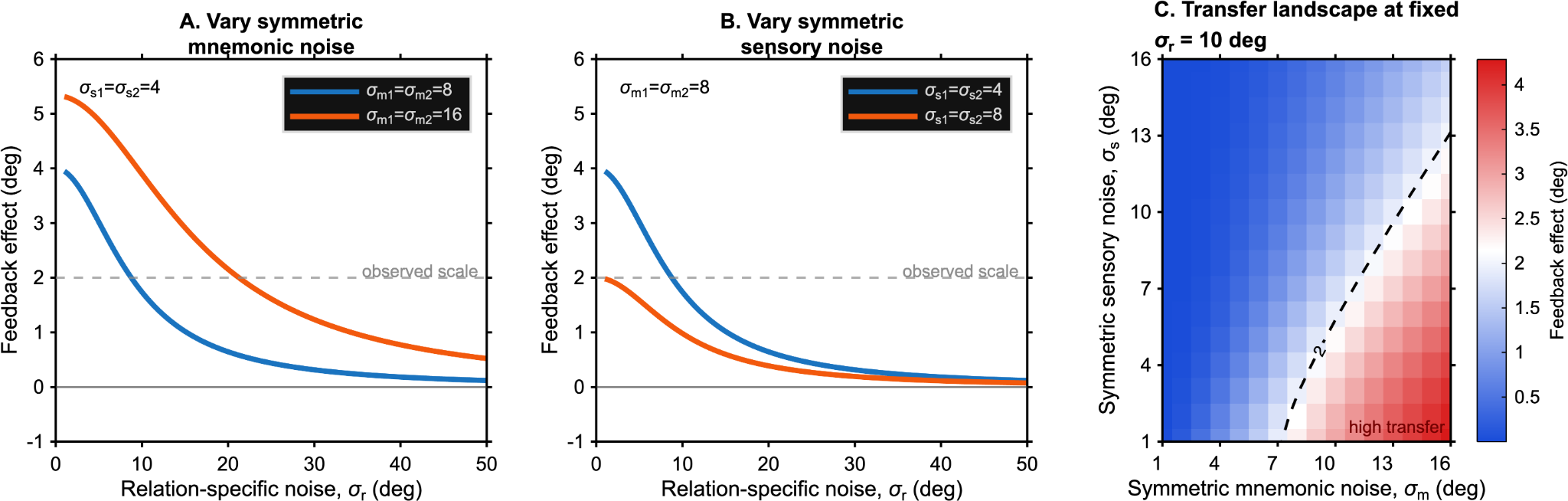

**Figure S5. The relation-constrained model predicts attenuated feedback transfer and distinct effects of mnemonic, sensory-stage, and relation-specific uncertainty.** Analytic predictions were generated from the posterior-conditioning gain of the relation-constrained model, in which feedback is modeled by conditioning one latent orientation on the feedback-specified value rather than by replacing its mnemonic evidence. Feedback transfer was quantified as the positive-minus-negative feedback contrast for the orientation for which feedback was not provided. Because the feedback values were displaced by $3^{\circ}$in opposite directions, complete transfer corresponds approximately to a $6^{\circ}$contrast under the assumptions described in the text; the gray dashed line indicates a contrast of approximately $2^{\circ}$, similar in scale to that observed in the experiments. **(A)** Predicted feedback transfer as a function of relation-specific noise, $\sigma_{r}$, for two levels of symmetric item-specific mnemonic noise, with $\sigma_{s1}=\sigma_{s2}=4^{\circ}$. Transfer decreases as relation-specific noise increases but is larger when item-specific mnemonic noise is higher, because the other orientation is less strongly anchored by its own mnemonic evidence and is therefore more influenced by the relation-based prediction. **(B)** Predicted feedback transfer as a function of $\sigma_{r}$for two levels of symmetric sensory-stage noise, with $\sigma_{m1}=\sigma_{m2}=8^{\circ}$. Transfer is weaker when sensory-stage noise is higher because increased marginal uncertainty reduces the posterior-conditioning gain. **(C)** Feedback-transfer landscape at fixed relation-specific noise, $\sigma_{r}={10}^{\circ}$, across symmetric sensory-stage and item-specific mnemonic noise values ranging from $1^{\circ}$to ${16}^{\circ}$. Warmer colors indicate stronger transfer. The dashed contour marks an approximately $2^{\circ}$feedback contrast. The strongest transfer is predicted when sensory-stage noise is low and item-specific mnemonic noise is high.

##### S3.5 Detectability of relation-preserving changes in joint posterior space

Although the present formulation was developed to explain feedback-induced transfer, the same joint-posterior geometry also yields a broader implication for change detection in multi-item memory. Previous studies have shown that observers can be relatively insensitive to changes that preserve relational or grouping structure, while being more sensitive to changes that disrupt such structure (Jiang et al., 2000; Orhan & Jacobs, 2013; Bateman et al., 2018; see also Chen & Levi, 1996; Lew & Vul, 2015). The present framework provides a possible probabilistic account of this pattern by asking how plausible different probe states are under the remembered joint posterior.

The model makes a relative prediction for changes that are matched in their item-wise deviations from the remembered state. When the absolute deviations of the individual item values are held constant, a relation-preserving change avoids the additional penalty associated with distorting the remembered offset, whereas a relation-distorting change incurs this relational penalty. Thus, relation-preserving changes are predicted to be relatively harder to detect than relation-distorting changes when item-wise deviation is matched.

Within the local Gaussian approximation used above, let the remembered posterior under model $M$ be

$$p_{M}\left( \theta\mid y_{M} \right)\mathcal{=N}\left( \mu_{M|y},\Sigma_{M\mid y} \right),$$

where

$$\theta=\left[ \begin{matrix} \theta_{1} \\ \theta_{2} \end{matrix} \right], \mu_{M|y}=\left[ \begin{matrix} \mu_{1} \\ \mu_{2} \end{matrix} \right].$$

Let the probe-defined item values be

$$\theta^{P}=\left[ \begin{matrix} \theta_{1}^{P} \\ \theta_{2}^{P} \end{matrix} \right],$$

with probe-defined offset

$$\Delta^{P}=\theta_{2}^{P}-\theta_{1}^{P}.$$

In the limiting case of negligible probe noise, evaluating the plausibility of the probe reduces to evaluating the remembered posterior density assigned to this probe state:

$$p_{M}\left( \theta^{P}\mid y_{M} \right)\propto exp\left[ -\frac{1}{2}D_{M}^{2}\left( \theta^{P} \right) \right],$$

where

$$D_{M}^{2}\left( \theta^{P} \right)=\left( \theta^{P}-\mu_{M} \right)^{\top}\Sigma_{M\mid y}^{-1}\left( \theta^{P}-\mu_{M} \right)$$

is the squared Mahalanobis distance of the probe from the center of the remembered posterior. A probe in a lower-probability region has a larger value of $D_{M}^{2}$ and is therefore more likely to be judged as changed, regardless of the precise decision criterion used downstream.

If probe noise is included, the same idea can be expressed with a posterior predictive distribution. Independent isotropic probe noise broadens the distribution without changing its principal directions; it may reduce the magnitude of the contrast but preserves the geometric ordering derived below.

##### Independent-item model

In the independent-item model, if the two item uncertainties are matched, the posterior covariance is diagonal and isotropic:

$$\Sigma_{I\mid y}=VI_{2}.$$

Consider two probe perturbations from the remembered posterior mean. A relation-preserving perturbation moves both item values by the same amount:

$$\theta_{\mathrm{pres}}^{P}=\mu_{I|y}+\left[ \begin{matrix} c \\ c \end{matrix} \right].$$

A relation-distorting perturbation is chosen to have the same absolute item-wise displacement, but to change the offset between the two items:

$$\theta_{\mathrm{dist}}^{P}=\mu_{I|y}+\left[ \begin{matrix} -c \\ c \end{matrix} \right].$$

The two probes therefore have the same Euclidean distance from the posterior mean and the sme absolute item-wise displacement (**Fig. S6, left**). In the independent-item model, their squared Mahalanobis distances are also identical:

$$D_{I}^{2}\left( \theta_{\mathrm{pres}}^{P} \right)=\frac{2c^{2}}{V}, D_{I}^{2}\left( \theta_{\mathrm{dist}}^{P} \right)=\frac{2c^{2}}{V}.$$

Thus, when the amount of item-wise deviation is matched, the independent-item model provides no basis for distinguishing the plausibility of relation-preserving from relation-distorting probes.

##### Relation-constrained model

In the symmetric relation-constrained model,

$$\Sigma_{R\mid y}=\left[ \begin{matrix} V & C \\ C & V \end{matrix} \right], C>0.$$

As shown in Section S3.2, the posterior variances along the normalized common and difference directions are

$$V_{\mathrm{common}}=V+C$$

and

$$V_{\mathrm{difference}}=V-C,$$

respectively. **The relation-preserving perturbation** ${[c,c]}^{T}$**lies along the common direction, whereas the relation-distorting perturbation** ${[-c,c]}^{T}$ **lies along the difference direction.** Their Mahalanobis distances are therefore

$$D_{R}^{2}\left( \boldsymbol{\theta}_{\mathrm{pres}}^{P} \right)=\frac{2c^{2}}{V+C}=\frac{2c^{2}}{V_{\mathrm{common}}},$$

and

$$D_{R}^{2}\left( \boldsymbol{\theta}_{\mathrm{dist}}^{P} \right)=\frac{2c^{2}}{V-C}=\frac{2c^{2}}{V_{\mathrm{difference}}}.$$

Because relational evidence constrains the difference direction more strongly,

$$V_{\mathrm{difference}}<V_{\mathrm{common}},$$

and therefore

$$D_{R}^{2}\left( \theta_{\mathrm{pres}}^{P} \right)<D_{R}^{2}\left( \theta_{\mathrm{dist}}^{P} \right).$$

Thus, the relation-preserving probe remains in a higher-density region of the remembered joint posterior, whereas the relation-distorting probe falls into a lower-density region (**Fig. S6, right**). **The difference does not arise because relational information is assumed to dominate item-specific information. Rather, once item-wise displacement is matched, the relation-distorting probe additionally moves against the more tightly constrained difference direction of the posterior.**

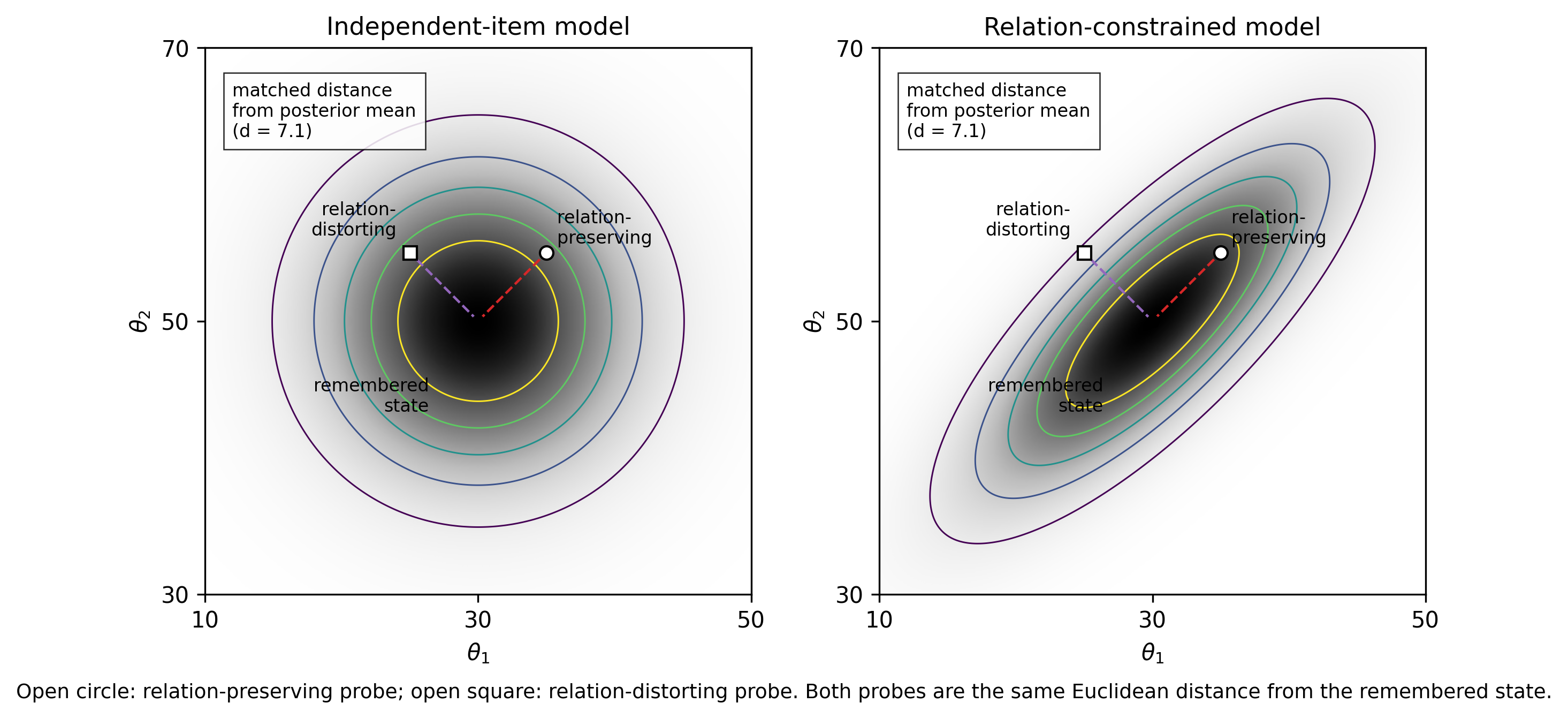

**Figure S6. Schematic illustration of relation-preserving and relation-distorting probe states in joint posterior space.** In each panel, the black point indicates the posterior mean, and the two open markers indicate probe states matched in Euclidean distance from that mean and in the absolute displacement of each item value. In the independent-item model, the posterior is isotropic in this schematic example, so the relation-preserving and relation-distorting probes fall on the same posterior contour and are equally plausible. In the relation-constrained model, relational evidence elongates the joint posterior along the direction in which the remembered offset is preserved. The relation-preserving probe lies along this higher-variance direction and remains relatively plausible, whereas the relation-distorting probe moves along the more tightly constrained difference direction and falls into a lower-density region. This illustrates why, holding item-wise displacement constant, relation-preserving changes are predicted to be relatively harder to detect than relation-distorting changes.

**The exact comparison outside the symmetric case depends on the full posterior covariance and on the directions of the probe perturbations. Nevertheless, the same qualitative prediction holds whenever relational evidence leaves greater posterior variance along a relation-preserving direction than along a matched relation-distorting direction.** Under those conditions, perturbations that preserve the remembered relation remain more compatible with the remembered joint distribution, whereas perturbations that distort the relation move across its more tightly constrained structure.

Thus, the present account is not limited to feedback-induced transfer. More generally, it formalizes how relational evidence determines which combinations of item values are probable in memory. Relational information does not merely coexist with item-specific information; it reshapes the joint posterior over possible multi-item states. **The resulting posterior geometry provides a common probabilistic basis for feedback-induced transfer and for the predicted relative difficulty of detecting relation-preserving changes when item-wise deviations are matched. This change-detection consequence is a model implication rather than a result directly tested in the present experiments.**

**References**

Bateman, J. E., Ngiam, W. X., & Birney, D. P. (2018). Relational encoding of objects in working memory: Change detection performance is better for violations in object relations. *PLOS ONE, 13*(9), e0203848.

Chen, S., & Levi, D. M. (1996). Angle judgment: Is the whole the sum of its parts? *Vision Research, 36*(12), 1721–1735.

Jiang, Y., Olson, I. R., & Chun, M. M. (2000). Organization of visual short-term memory. *Journal of Experimental Psychology: Learning, Memory, and Cognition, 26*(3), 683–702.

Lew, T. F., & Vul, E. (2015). Ensemble clustering in visual working memory biases location memories and reduces the Weber noise of relative positions. *Journal of Vision, 15*(4), 10.

Orhan, A. E., & Jacobs, R. A. (2013). A probabilistic clustering theory of the organization of visual short-term memory. *Psychological Review, 120*(2), 297–328.
